# PCSK5^M452I^ is a recessive hypomorph exclusive to MCF10DCIS.com cells

**DOI:** 10.1101/2025.03.03.641323

**Authors:** Taylor Marohl, Kristen A. Atkins, Lixin Wang, Kevin A. Janes

## Abstract

The most widely used cell line for studying ductal carcinoma in situ (DCIS) premalignancy is the transformed breast epithelial cell line, MCF10DCIS.com. During its original clonal isolation and selection, MCF10DCIS.com acquired a heterozygous M452I mutation in the proprotein convertase PCSK5, which has never been reported in any human cancer. The mutation is noteworthy because PCSK5 matures GDF11, a TGFβ-superfamily ligand that suppresses progression of triple-negative breast cancer. We asked here whether PCSK5^M452I^ and its activity toward GDF11 might contribute to the unique properties of MCF10DCIS.com. Using an optimized in-cell GDF11 maturation assay, we found that overexpressed PCSK5^M452I^ was measurably active but at a fraction of the wildtype enzyme. In a *PCSK5^−/−^* clone of MCF10DCIS.com reconstituted with different PCSK5 alleles, PCSK5^M452I^ was mildly defective in anterograde transport. However, the multicellular organization of PCSK5^M452I^ addback cells in 3D matrigel cultures was significantly less compact than wildtype and indistinguishable from a PCSK5^T288P^ null allele. Growth of intraductal MCF10DCIS.com xenografts was similarly impaired along with the frequency of comedo necrosis and stromal activation. In no setting did PCSK5^M452I^ exhibit gain-of-function activity, leading us to conclude that it is hypomorphic and thus compensated by the remaining wildtype allele in MCF10DCIS.com.

**Implications:** This work reassures that an exotic PCSK5 mutation is not responsible for the salient characteristics of the MCF10DCIS.com cell line.

## INTRODUCTION

Ductal carcinoma *in situ* (DCIS) is a prevalent yet enigmatic premalignancy of the breast with a ∼30% chance of progressing to invasive cancer (1). DCIS latency is thought to be long and variable in women, and its molecular subtypes differ from those of breast carcinoma (2,3). The biology of DCIS is notoriously difficult to access experimentally. There are long-standing genetically-engineered mouse models of breast cancer that have well defined DCIS intermediates. However, they are driven by viral transgenes [polyomavirus middle T antigen (4), SV40 large T antigen (5)] and progress rapidly without the indolence characteristic of human DCIS (6). This prompted early efforts to derive human cell lines that capture facets of DCIS *in vitro* and *in vivo*.

Nearly 25 years ago, Miller *et al*. described MCF10DCIS.com as a derivative of MCF10A cells (7) that was transformed with oncogenic HRAS^G12V^ (8), subcloned, and propagated subcutaneously in nude mice (9). MCF10DCIS.com is singular among HER2-negative cell lines in initiating durable xenografts with comedo necrosis that are histologically DCIS-like (10). It is a pillar of the MCF10 series of cell lines (11) and has served as a reference for DCIS research decades before the exciting recent advances with intraductal xenotransplantation of patient-derived samples (12). MCF10DCIS.com cells underlay impactful studies of malignancy (10), metastasis (13), metabolism (14,15), and mechanotransduction (16). The line remains useful for spanning complexity scales—MCF10DCIS.com expands readily in 2D tissue culture and 3D matrigel culture, as well as in the subcutaneous tissue, mammary duct, and mammary fat pad of immunocompromised mice (17–21). Although there are valid concerns about the initiating oncogene (22), MCF10DCIS.com cells are generally viewed as a best-available proxy for human hormone receptor-negative DCIS.

MCF10DCIS.com cells harbor ectopic HRAS^G12V^ (8), an acquired PIK3CA^H1047R^ driver mutation (15), as well as additional somatic changes of unknown significance (23). Among the most unusual is a M452I point substitution in the proprotein convertase, PCSK5 (24). Mutant *PCSK5* is detectable in transformed MCF10 predecessors and increases to 50% allele frequency in MCF10DCIS.com before receding in MCF10 invasive carcinoma lines; no other mutation exhibits this pattern (23). The most-recognized convertase substrate for PCSK5 is GDF11 (25,26), a secreted TGFβ superfamily ligand that modulates cancer progression (27). In breast cancer, we previously showed that GDF11 promotes epithelial organization, restrains invasion, and loses activity when its inactive precursor is not matured due to PCSK5 silencing or mutation (28). Multiple breast cancer-derived PCSK5 mutants were validated as loss-of-function in our earlier study. However, M452I is not documented in breast cancer nor in any cancer to date (29), and a loss-of-function allele would be counterintuitive given the anti-invasive characteristics of MCF10DCIS.com. More plausible was that PCSK5^M452I^ conferred a non-canonical gain-of-function phenotype and matured endogenous proGDF11 more efficiently than wildtype, as recently demonstrated for a V474I polymorphism in a related proprotein convertase (30). If true, it would provide a satisfying explanation for the select characteristics of MCF10DCIS.com and call into question prior conclusions that might have stemmed from a one-off mutation never before documented in human disease.

Here, we performed a detailed investigation of the PCSK5^M452I^ mutant derived from MCF10DCIS.com. Using an orthogonal mammalian-expression system, we designed a proGDF11 convertase assay, which suggested that PCSK5^M452I^ was detectably less active than wildtype PCSK5, with additive effects when the two were coexpressed. MCF10DCIS.com-specific properties were examined by knocking out PCSK5 and inducibly reconstituting with PCSK5, PCSK5^M452I^, or a known loss-of-function allele [PCSK5^T288P^ (28)]. In these cells, we quantified subcellular protein localization in 2D monolayers, multicellular organization in 3D cultures, and *in vivo* histopathology of intraductal inocula. The PCSK5^T288P^ addback cells verified PCSK5 loss-of-function and phenocopied prior results involving GDF11 knockdown (28), but in no context was there a hypermorphic or neomorphic difference between wildtype PCSK5 and PCSK5^M452I^ addback. Our reassuring conclusion is that PCSK5^M452I^ is a hypomorphic allele, which is heterozygous recessive and thus unlikely to drive phenotypes in MCF10DCIS.com.

## MATERIALS AND METHODS

### Cell lines

MCF10DCIS.com cells were obtained directly from the Karmanos Cancer Institute through a Material Transfer Agreement, and cell identity was confirmed by STR profiling. MCF10DCIS.com cells were maintained in DMEM/F-12 medium (Gibco, 11320) plus 5% horse serum (Gibco, 16050) and 3D cultured in MCF10A assay medium (31). MCF10A-5E cells were isolated and maintained as previously described (32). HEK 293T/17 cells (ATCC, CRL-11268) were cultured in DMEM (Gibco, 11965) plus 10% fetal bovine serum (HyClone, SH303396.03). All base media were supplemented with 1% penicillin–streptomycin (Gibco, 15140), and all cell lines were cultured at 37°C.

### Plasmids

GDF11 secretion assay—pLX302 GDF11-V5 puro (Addgene, 83097), pLX304 (wildtype) PCSK5-V5 blast (Addgene, 83100), and pLX304 PCSK5 (T288P)-V5 blast (Addgene, 83101) were described previously (28). pDONR223 PCSK5 (M452I) (Addgene, 232445) was prepared by QuikChange II XL site-directed mutagenesis (Agilent, 200521) of pDONR223 (wildtype) PCSK5 from the human ORFeome v5.1 and recombined into pLX304 (Addgene, 25890) with LR clonase II (Invitrogen, 11791020) to yield pLX304 PCSK5 (M452I)-V5 blast (Addgene, 232446). pcDNA3 was used as a carrier plasmid for lipofections, and pLX302 EGFP-V5 puro (Addgene, 141348) or pLX304 EGFP-V5 blast (Addgene, 232447) was used when diluting GDF11 or PCSK5 plasmid dosage and for negative controls. pcDNA3.1 HRAS (G12V) was kindly provided by David Kashatus.

Knockout and addback of PCSK5 alleles—For PCSK5 knockout, an sgRNA sequence (sg09, CTACACGGGAAAGAACATTG) was cloned into EDCPV (Addgene, 90085) by conventional restriction digest, oligo annealing, and ligation to yield EDCPV PCSK5_sg09 (Addgene, 232455). For PCSK5 addback, an inducible donor plasmid for PCSK5 was first constructed by PCR cloning of wildtype PCSK5 with a C-terminal V5 epitope tag into the SpeI–MfeI sites of pEN_TTmiRc2 (Addgene, 25752) to yield pEN_TT PCSK5-V5 (Addgene, 232448). Next, an sgRNA-resistant (sgRR) allele of PCSK5 was prepared by introducing 3 silent mutations into pEN_TT PCSK5-V5 by QuikChange II XL site-directed mutagenesis (Agilent, 200521) to yield pEN_TT PCSK5(sgRR)-V5 (Addgene, 232449). The sgRR plasmid was further mutagenized to yield pEN_TT PCSK5(sgRR,M452I)-V5 (Addgene, 232450) and pEN_TT PCSK5(sgRR,T288P)-V5 (Addgene, 232451). Last, the three sgRR donor plasmids were recombined into pSLIK hygro (Addgene, 25737) with LR clonase II (Invitrogen, 11791020) to yield pSLIK PCSK5(sgRR)-V5 hygro (Addgene, 232452), pSLIK PCSK5(sgRR,M452I)-V5 hygro (Addgene, 232453), and pSLIK PCSK5(sgRR,T288P)-V5 hygro (Addgene, 232454).

Bioluminescence imaging—We removed the V5 epitope tag of pLenti PGK Blast V5-LUC (w528-1) (Addgene, 19166) by BamHI digestion and self-ligation to yield pLenti PGK Blast LUC (w528-1) (Addgene, deposition pending).

### Computational assessment of PCSK5 mutations

Computational predictions for the mutations R486H (rs138257548, chr9:76159009G>A), A565T (rs145509473, chr9:76169777G>A), M452I (chr9:76157088G>T), and T288P (COSV65097476, chr9:76071866A>C) were made in November 2024 with PCSK5A (transcript ID NM_006200.6/ENST00000376752.9, protein ID NP_006191.2/ENSP00000365943.4). For AlphaMissense (33,34), PCSK5A was not available, and so PCSK5B (transcript ID ENST00000545128.5) was used instead. Predictions using AlphaMissense (33,34), Cscape (35), MutationTaster2021 (36), and PANTHER-PSEP (37) were taken directly from the corresponding website with no change in default parameters. Predictions using FATHMM v2.3 (38), LRT (39), Meta-RNN (40), MutationAssessor r3 (RRID: SCR_024502), SIFT (41), and SIFT 4G 2.4 (42) were aggregated by dbNSFP v4.7 (43). Predictions from computational tools were categorized as neutral (N), likely neutral (LN), unknown (U), likely damaging (LD), or damaging (D) as described in Supplementary Table S1.

### *PCSK5* genotyping

Genomic DNA was isolated from standard cultures of MCF10DCIS.com and MCF10A-5E cells with the DNeasy Blood & Tissue Kit (Qiagen, 69504). The *PCSK5* locus was PCR amplified and sequenced with the following primers: GTGGGGCCCTGGAGAAAAA (forward), AAGAGCCCAGGGGTAAGCAT (reverse), ATCCCATAGGGTGGTGTCTG (sequencing). Total RNA was isolated from MCF10DCIS.com cells with the RNeasy Mini Kit (Qiagen, 74134) after culturing cells for 6 days in assay medium (31) plus 2% growth factor-reduced matrigel (Corning, 356253) and 5 ng/ml EGF (Peprotech, AF-100-15). Total RNA was similarly isolated from MCF10A-5E cells cultured in growth medium. 250 ng RNA was reverse transcribed with oligo(dT)_24_ and Superscript III (Invitrogen, 18080085), and the transcribed *PCSK5* locus was PCR amplified and sequenced with the following primers: CCCTGCCAGTCTGACATGAA (forward), TTTGTCGGTCTGTGCTCTCC (reverse), GTACCTGGAAGAGTGTTCATCC (sequencing).

### PCSK5 antibody

The custom PCSK5 antibody was raised in rabbit to target the peptide sequence Ac-SPTNEFPKVERFRYSRC-amide (PCSK5-A amino acids 604–619) and affinity purified to a stock concentration of 1.82 mg/ml (Covance).

### Quantitative immunoblotting

Cells were lysed in RIPA buffer supplemented with proteinase inhibitor cocktail, except for whole-cell PCSK5-V5 quantification, where cells were trypsinized, counted, and lysed in Laemmli sample buffer. Quantitative immunoblotting was performed as described (44) with primary antibodies recognizing the following targets: CDH1 (BD Biosciences, 610182; 1:1000 dilution), ERK1/2 (EMD Millipore, ABS44; 1:2000 dilution), HRAS (Santa Cruz, sc-520; 1:1000 dilution), p38 (Santa Cruz, sc-535; 1:5000 dilution), PCSK5 (Covance, custom; 1:1000 dilution), TP63 (Biocare, 163A;, 1:500 dilution), tubulin (chicken polyclonal—Abcam, ab89984; 1:20,000 dilution or rabbit polyclonal—Cell Signaling, 2148; 1:2000 dilution), V5 (mouse monoclonal—Invitrogen, 46-0705; 1:5000 dilution or chicken polyclonal—Bethyl Laboratories, A190-118A; 1:5000 dilution), VIM (Abcam, ab16700; 1:300 dilution), and vinculin (Millipore, 05-386; 1:10,000 dilution). Primary antibodies were probed with IRDye 680RD-or 800CW-conjugated secondary antibodies (LI-COR; 1:20,000 dilution) and visualized on a LI-COR Odyssey infrared imaging system. For endogenous PCSK5 detection, a tertiary detection scheme was used involving unconjugated AffiniPure goat anti-rabbit IgG (Jackson ImmunoResearch,111-005-144; 1:1000 dilution) followed by IRDye 800CW-conjugated donkey anti-goat IgG (LI-COR, 926-32214; 1:20,000 dilution). For whole-cell PCSK5-V5 quantification, calibration was performed with serial dilutions of V5-containing recombinant Multitag protein (GenScript, M0101).

### ProGDF11 convertase assay

HEK 293T/17 cells (ATCC, CRL-11268) were seeded in 24-well plates at a density of 100,000 cells per cm^2^ one day before lipofection with Lipofectamine 3000 (Invitrogen, L3000015) and 500 total ng of pLX304 PCSK5-V5 blast (0–40 ng, with 3 ng optimal), pLX302 GDF11-V5 puro (100 ng), and pcDNA3 carrier plasmid (400–360 ng). Existing medium was removed before gentle addition of lipocomplexes in 100 µl DMEM (Gibco, 11965) combined with 250 µl growth medium per well. Edge wells were left unplated and filled with PBS to minimize evaporation effects of lipofected cells. After 24 hours, conditioned medium was collected, and the concentration of secreted GDF11 was quantified with a GDF11 ELISA (R&D Systems, DY1958). If needed, samples were pre-diluted in Reagent Diluent (R&D Systems, DY008B) to remain within the standard range of the assay (31–2000 pg/mL). The calibration curve was modeled by four parameter logistic regression and used to calculate concentration of mature GDF11 in unknown samples. When PCSK5 or GDF11 were reduced or omitted as controls, the corresponding EGFP plasmid was substituted. For HRAS^G12V^ experiments, cells were co-transfected with a PCSK5 allele, GDF11, and 2 ng pcDNA3 HRAS^G12V^. All secreted GDF11 concentrations were normalized to the GDF11/EGFP control condition from the same experiment.

### Immunoprecipitation

GDF11-V5 was immunoprecipitated from 250 µl conditioned medium of transfected HEK 293T/17 cells with 1 µg mouse anti-V5 (Invitrogen, 46-0705) overnight at 4°C followed by 10 µl Protein A/G Plus Ultralink Resin beads (Thermo Fisher, 53135) for one hour at 4°C, followed by two washes with ice-cold PBS. Beads were boiled in 2x Laemmli sample buffer and immunoblotted with chicken anti-V5 (Bethyl Laboratories, A190-118A; 1:5000 dilution).

### Lentiviral transduction and cell selection

MCF10DCIS.com cells were seeded in 6-well plates at a density of 50,000 cells per well one day before three serial daily transductions with 500 µl of EDCPV PCSK5_sg09 lentivirus freshly prepared by calcium phosphate transfection of HEK 293T/17 cells (ATCC, CRL-11268) (45). Transductants were treated with 200 nM Shield-1 (AOBIOUS, AOB1848) in MCF10A assay medium supplemented with 5 ng/ml EGF (Peprotech, AF-100-15) and 2% growth factor-reduced matrigel (Corning, 356253) (31) for 12 days. Venus-expressing cells were clonally sorted into 96-well plates with a Sony MA900 Cell Sorter (Sony MA900) at the University of Virginia Flow Cytometry Core and supplemented with 10 ng/ml recombinant GDF11 (Peprotech, 120-11) during clonal expansion. Genomic DNA was harvested from clones with the PureLink Genomic DNA Mini Kit (Thermo Fisher, K182001) and the PCSK5-sg09 target site was PCR-amplified (Millipore Sigma, 11732650001) with primers CAGCATGCTCTTCTTCTTTCAG (forward) and GAGTGTATGCTGTGGTTAGAAGGTC (reverse). Amplicons were cloned into TOPO plasmids (Invitrogen, 450030) and Sanger sequenced to verify successful PCSK5 knockout. GDF11 supplementation was removed after knockout verification to facilitate the subsequent lentiviral transduction. MCF10DCIS.com PCSK5^−/−^ cells were transduced with 100 µl of pSLIK PCSK5(sgRR)-V5 hygro, pSLIK PCSK5(sgRR,M452I)-V5 hygro, or pSLIK PCSK5(sgRR,T288P)-V5 hygro lentivirus prepared by calcium phosphate transfection of HEK 293T/17 cells and stored at –80°C (45). Transductants were selected with 100 µg/ml hygromycin (Sigma, H0654) until control plates had cleared.

### Immunofluorescence

PCSK5 addback cells were seeded on coverslips in 6-well plates (275,000 cells per well) in MCF10DCIS.com growth medium supplemented with 100 µg/ml hygromycin (Sigma, H0654) and 1 µg/ml doxycycline. After approximately 24 hours, cells were fixed in ice-cold methanol for 5 minutes at –20°C, and then immunofluorescence was performed as previously described (45) for the following targets: PDI (Thermo Fisher, MA3-019, 1:200 dilution), GM130 (BD Biosciences, 610823; 1:750 dilution), TGN38 (Thermo Fisher, MA3-063; 1:200 dilution), V5 (Bethyl, A190-118A; 1:1000 dilution), and α/ꞵ-Tubulin (Cell Signaling Technology, 2148; 1:100 dilution). Primary antibodies were visualized with Alexa Fluor 488-conjugated goat anti-mouse (Invitrogen, A11029; 1:200 dilution), Alexa Fluor 555-conjugated goat anti-chicken (Invitrogen, A21437; 1:200 dilution), and Alexa Fluor 647-conjugated goat anti-rabbit (Invitrogen, A21245; 1:200 dilution). After counterstaining with DAPI, coverslips were incubated with 10 mM CuSO_4_ in 50 mM NH4Ac (pH 5.0) for 10 minutes at room temperature (46) and then washed once with PBS before mounting. Mounted coverslips were imaged as 5–13 optical sections on a Leica STELLARIS 5 LIAchroic confocal laser-scanning microscope with a 63x 1.4 NA plan apochromat oil-immersion objective and the following acquisition parameters: 70.7 nm pixel size at 1.28x optical zoom; 2048 x 2048 pixels^2^ field of view; 167 ns pixel dwell time; 500 nm z step size; 1 Airy Unit pinhole for a 520 nm emission; line accuracy = 2; 405 nm laser power = 2%; 488 nm laser power = 0.2% (PDI), 0.5% (GM130), or 2% (GM130, TGN38); 561 nm laser power = 2%; 638 nm laser power = 5%; and all detectors in photon-counting mode. Image stacks were deconvolved with Leica LIGHTNING deconvolution software using default parameters and an immersion medium of 90% glycerol + 10% water.

### Fluorescence segmentation and image analysis

We used CellProfiler (47) version 4.2.6 to analyze the optical section in each image stack that best captured secretory pathway staining for most cells. Each multicolor image was first thresholded above a photon count of 10 (V5, PDI, GM130, and TGN38) or 5 (DAPI). The DAPI signal was converted to a nuclear mask by a series of morphological operations (fill, diagonal, bridge, majority) to fill holes. The nuclear mask was globally thresholded with a diameter range of 80–350 pixels and then shrunk by 4 pixels. The tubulin signal was smoothed and combined with the nuclear mask to create a cell mask that was subsequently expanded by 10 pixels to ensure complete capture of the cell periphery. Colocalization of V5 with PDI, GM130, and TGN38 in the whole-cell mask of each cell was quantified by the Manders colocalization coefficient relative to total V5 immunoreactivity after excluding pixels with photon counts below the specified threshold for each marker. Colocalization coefficients were arcsine-transformed and batch corrected by study to account for variability in the extent of PCSK5 induction between days. The batch correction was hierarchically defined as the global mean of the arcsine-transformed means of each V5-organelle stain for that day. Mean V5 intensity in the nuclear mask of each cell was log-transformed and batch corrected by study. Cells with no above-threshold organelle staining in the optical section (0.03% of all cells segmented) were excluded from the analysis.

### 3D matrigel culture

Eight-well chamber slides (Fisher, 354108) were coated with 40 µl growth factor-reduced matrigel (Corning, 356253) and centrifuged at 4°C for 10 minutes at 1824 rcf. MCF10DCIS.com cells were seeded at 5000 cells per well in 400 µl MCF10A assay medium supplemented with 5 ng/ml EGF and 2% matrigel as described (31). Where indicated, assay medium was supplemented with 250 ng/ml recombinant mature GDF11 protein (Peprotech, 120-11) throughout the experiment, or supplemented with 1 µg/ml doxycycline from Day 4 onward. Medium was replaced every 4 days, and cultures were imaged as 3×3 fields with a 4x apochromat air objective on an EVOS M7000 (Thermo Fisher, AMF7000) using DiamondScope software (version 2.0.2094.0).

### Brightfield segmentation and image analysis

Spheroids were segmented with an updated OrganoSeg (48) software kindly provided by Cameron Wells. All studies used the following parameter settings: out of focus correction = yes, DIC correction = no, split whole image = no, edge correction = yes, size threshold = 650, window size = 300, contaminant intensity = zero, minimum circularity = zero, and segmentation closing structuring element size = 2. Study-specific parameters that depended on overall illumination and focusing were: intensity threshold = 0.5–0.9, edge correction = 0.25–0.35, image reconstruction structuring element size = 2–3, and use reconstructed image for edge correction = true or false. All other parameters were set to the software default. Segments were manually excluded as contaminants if they captured obvious debris, bubbles, 2D growth, or background illumination artifacts; or, if the segment did not accurately reflect an out-of-focus spheroid. Other study-specific refinements included exclusion of partially segmented spheroids, removal of out-of-focus image fields, and splitting of interconnected spheroids when present. From retained image segments, we exported cross-sectional area and circularity for analysis. Area measurements were batch-corrected for experimentalist by normalizing to the global Day 4 mean area calculated after subtracting the segmentation size threshold (650 pixels^2^), adding an offset of 10 pixels^2^, and log transforming (49). When necessary, circularities were batch-corrected for experimentalist by normalizing to the global mean circularity after arcsine transformation.

### PCSK5 nuclear localization sequence prediction

Nuclear localization sequence predictions for PCSK5A (protein ID NP_006191) were made in March 2025 using cNLS Mapper (50) with a cutoff score of 2.0 or greater. The entire sequence was searched for bipartite nuclear localization sequences with a long linker.

### Intraductal inoculation

For inoculation, PCSK5^−/−^ MCF10DCIS.com cells with PCSK5 addback were transduced with pLenti PGK Blast LUC (w528-1) and selected with 10 µg/ml blasticidin until control plates had cleared. Female, virgin SCID/beige mice (Charles River, 250) were obtained at 7 weeks of age and housed together on a 10-hour dark cycle (8 pm–6 am) and 14-hour light cycle (6 am–8 pm) at 21.5°C and 31.5% relative humidity. Mice were given a standard rodent diet. Mice were anesthetized with isoflurane, depilated, and the nipple tip of the fourth mammary glands were cut off with a fine scissor. Each luciferase-expressing PCSK5 addback variant was suspended at a concentration of 20,000 cells/ml in growth medium, loaded into a 30-gauge blunt needle (Hamilton, 80508), and injected into the fourth and ninth mammary gland (2 µl per gland). Mouse surgical order and left-right gland assignments for the PCSK5 genotypes (*N* = 10 injections per genotype) were randomized. Mice were switched to rodent food containing 625 mg/kg doxycycline (Harlan, TD.05125, IF060, HF030) starting one week after surgery, which was maintained until the end of the study. All animal work was done in compliance with ethical regulations under University of Virginia IACUC approval #3945, which permitted a maximal tumor size of 2.0 cm that was not exceeded by this study.

### Bioluminescent imaging and analysis

Mice were imaged for bioluminescence at 2, 7, 14, 21, 28, 35, 42, 49, and 54 days post-surgery after isoflurane anesthesia and intraperitoneal administration of 150 µg D-luciferin (Promega, E1605) per gram body weight. At 5–10 minutes post-injection of D-luciferin, bioluminescence was collected every 2 minutes for 15–20 minutes on a Lago X (Spectral Instruments Imaging) with Aura Imaging Software (version 4.0.7) at the University of Virginia Molecular Imaging Core. The peak emission (photons/s) across all 2-minute images per gland was taken as the bioluminescence readout for that time point after inoculation. Bioluminescence was compared longitudinally by normalizing each gland to its peak emission averaged across Days 2, 7, and 14. This normalization accounts for gland-to-gland differences in inoculum volume and viability. Multiway ANOVAs considered cage number and left-right gland injections as fixed effects alongside PCSK5 genotype.

### Mammary gland harvest and histology

Mice were euthanized at 54 days post-surgery and perfused with 4% paraformaldehyde before gland harvesting (51) and additional fixation in 4% paraformaldehyde for 24 hours. Samples were paraffin embedded, cut as 4–5 µm sections, and hematoxylin–eosin stained by the University of Virginia Research Histology Core.

### Histological analysis

One or two representative sections from each mammary gland were selected for analysis after digital acquisition on an Aperio ScanScope. Sections were sub-divided into multiple ductal cross-sections with a distinct outer stroma. Ductal cross-sections were excluded from the analysis if (major axis) x (minor axis) ≤ 12,000 µm^2^, if the section was tangential to a duct, or if the minimum Feret diameter of the lesion was greater than 5 mm at dissection. Ductal cross-sections were scored positive or negative for the pathologic features summarized in Supplementary Table S2. Scores were summarized gland-wise as the fraction of ductal cross-sections positive for each feature.

### Statistics

All hypothesis tests are specified in the corresponding figure caption. Nonlinear least squares curve fitting was performed with the nlinfit function in MATLAB (Mathworks, R2023b), with confidence intervals estimated by the nlparci function.

## RESULTS

### The PCSK5^M542I^ allele expressed by MCF10DCIS.com is plausibly altered in function

In the first report of PCSK5^M452I^ (23), computational algorithms disagreed on the predicted severity of an M452I substitution in PCSK5. We revisited these inferences with newer methods (33,36,37,43,52) and evaluated PCSK5^M452I^ relative to two missense single-nucleotide polymorphisms (SNPs) in the short form of PCSK5—rs138257548 (encoding R486H) and rs145509473 (encoding A565T)—along with a somatic T288P mutation verified as inactive (28). Whereas PCSK5 SNPs were generally predicted to be neutral or minimally damaging, PCSK5^M452I^ was predicted to be damaging as frequently as PCSK5^T288P^ (**Fig. 1A**), warranting further study of PCSK5^M452I^ in MCF10DCIS.com.

**Figure 1.**
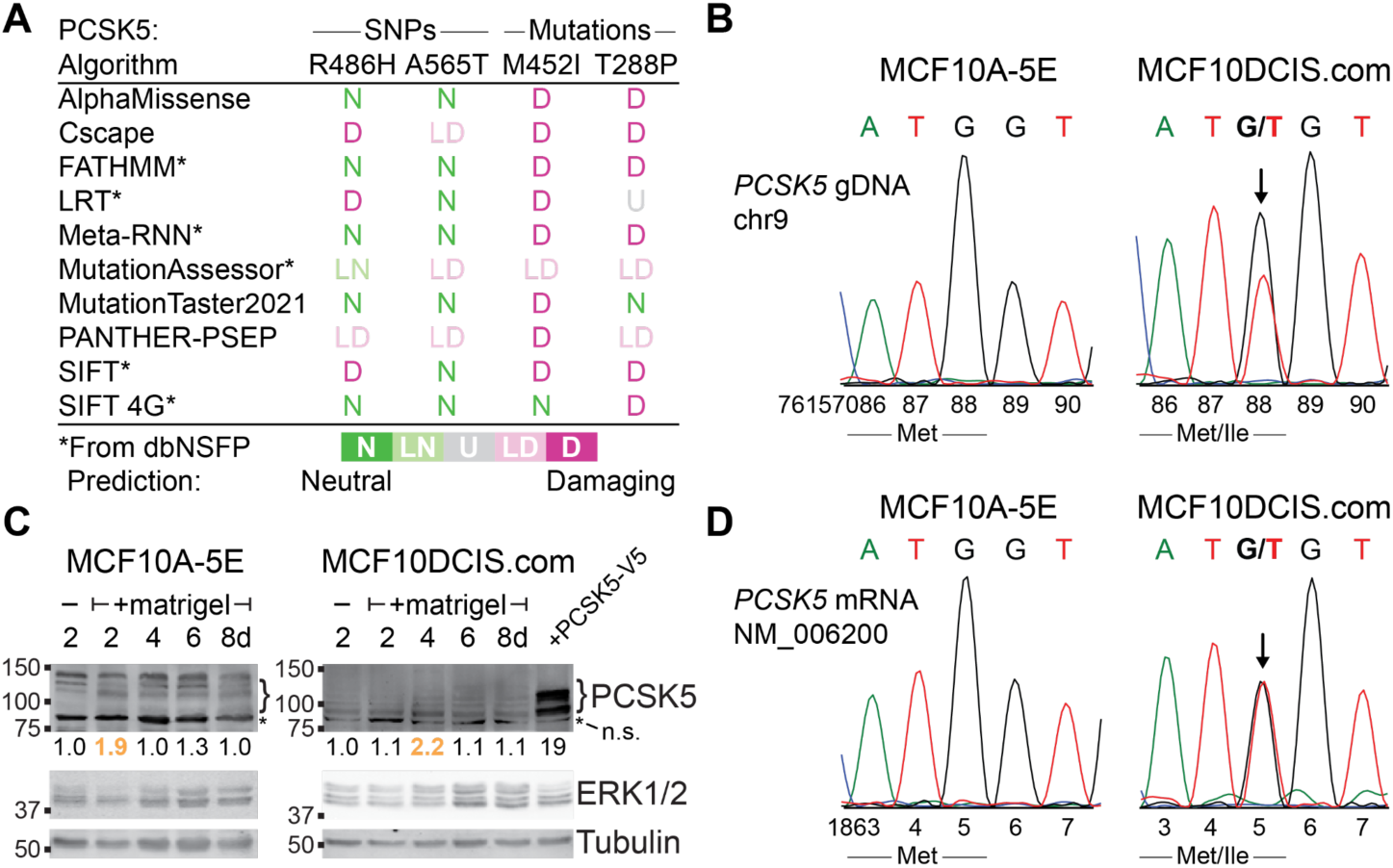
MCF10DCIS.com cells express a mutated allele of PCSK5 that is predicted to be damaging. **A,** Computational predictions for the PCSK5^M452I^ mutation of MCF10DCIS.com alongside two single-nucleotide polymorphisms (SNPs; R486H and A565T) and a somatic mutation (T288P) confirmed experimentally to be inactive (28). Outputs for each algorithm were grouped as Neutral (N), Likely Neutral (LN), Unknown (U), Likely Damaging (LD), or Damaging (D) as described in Supplementary Table S1. **B,** Confirmation of the M452I mutation in genomic DNA (gDNA) from MCF10DCIS.com cells. **C,** Endogenous PCSK5 is induced when MCF10-series cells are cultured with 2% matrigel for 2–4 days. Cell extracts were immunoblotted for PCSK5 with ERK1/2 and tubulin used as loading controls. Extract from MDA-MB-231 cells overexpressing pro and mature V5-tagged PCSK5 (+PCSK5-V5) was used as a positive control. Asterisk marks a nonspecific band for the PCSK5 antibody. Image gamma = 2 for the PCSK5 immunoblots. **D,** The PCSK5^M452I^ allele is expressed in MCF10DCIS.com cells cultured with 2% matrigel. For (**B**) and (**D**), MCF10A-5E cells (32) provide a wildtype reference.

We first confirmed the chr9:67157088G>T substitution in *PCSK5* at roughly 50% allele frequency (23) in genomic DNA prepared from MCF10DCIS.com, using the 5E clone (32) of MCF10A cells as a wildtype reference (**Fig. 1B**). To ensure both transcripts were equally abundant in MCF10DCIS.com, we required a context in which endogenous PCSK5 was reliably detected. However, *PCSK5* transcripts are very low in breast carcinomas and standard breast cancer cell lines (28), and commercial reagents for detecting PCSK5 protein were insensitive or nonspecific. We thus raised a new polyclonal antibody against residues 604–619, affinity purified the antisera, and optimized immunoblotting conditions to achieve fmol sensitivity against ectopic PCSK5 (Supplementary Fig. 1). Using the purified antibody, we discovered that PCSK5 is transiently induced when MCF10-series cells are cultured with 2% matrigel for multiple days (**Fig. 1C**). Total RNA extracted from matrigel-treated cells confirmed that both the wildtype and M452I transcripts were present in MCF10DCIS.com (**Fig. 1D**). These results supported that PCSK5^M452I^ protein was expressed when MCF10DCIS.com cells were exposed to certain microenvironments.

### An ectopic proGDF11 convertase assay suggests that PCSK5^M452I^ is enzymatically deficient

PCSK5 is the major convertase of proGDF11 (25,26). To quantify the activity of different PCSK5 alleles *in cellulo*, we developed a high-throughput GDF11 secretion assay by ectopically coexpressing PCSK5 and GDF11 in 293T cells and quantifying GDF11 release with a commercial ELISA. Endogenous GDF11 was barely detectable in conditioned medium from control lipofections and was generally low when GDF11 was overexpressed on its own (**Fig. 2A**, Lanes 1–5), confirming that endogenous convertase activity is modest in these cells (53). Cotransfection with PCSK5 hyperbolically increased the measured GDF11 ELISA signal, and we optimized wildtype PCSK5 gene dosage to provide dynamic range for increased activity as well as decreased activity (Supplementary Fig. 2A). In principle, the capture and detection antibodies of the ELISA might recognize unprocessed proGDF11 in the medium, which would obscure GDF11 maturation by conflating it with proGDF11 release. Using conditioned medium from optimized lipofections, we immunoprecipitated GDF11 by its epitope tag and immunoblotted to separate the pro and mature proteoforms by molecular weight. Consistent with the ELISA results, we observed much less mature GDF11 from cells that were not cotransfected with PCSK5 (Supplementary Fig. 2B). Across a range of conditions, the ELISA readout correlated with mature GDF11 abundance and anticorrelated with proGDF11 released into the medium (Supplementary Fig. 2C and D). These results indicated that the ELISA is specific for mature GDF11; thus, the secretion assay largely measures proGDF11 conversion by the PCSK5 that is cotransfected.

**Figure 2.**
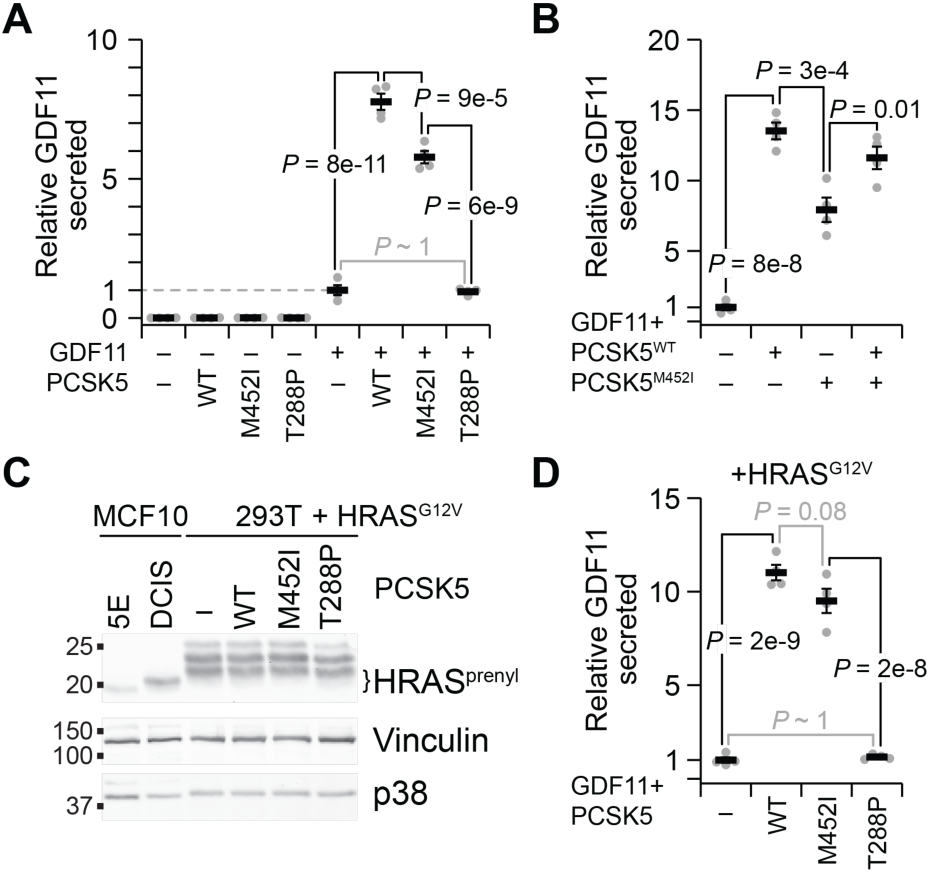
293T cells co-expressing PCSK5^M452I^ secrete mature ectopic GDF11 with intermediate efficiency. **A,** Catalytically active PCSK5 promotes ectopic GDF11 secretion. Cells were lipofected with 3 ng of the indicated PCSK5 allele (or EGFP overexpression control) plus 100 ng of GDF11, and conditioned medium was collected after 24 hours to measure GDF11 release by ELISA. PCSK5^T288P^ was included as a catalytically dead control (28). **B,** GDF11 release by PCSK5^M452I^ plus wildtype PCSK5 (PCSK5^WT^) is additive. Cells were treated as in (**A**) and compared with 1.5 ng PCSK5^WT^ plus 1.5 ng PCSK5^M452I^. **C,** Cotransfection with oncogenic HRAS^G12V^ approximates the level of prenylated HRAS (HRAS^prenyl^) in MCF10DCIS.com. 293T cells were lipofected with 2 ng of HRAS^G12V^ and 3 ng of the indicated PCSK5 allele (or EGFP overexpression control) and immunoblotted for HRAS with vinculin and p38 used as loading controls. MCF10A-5E cells are a negative control for HRAS overexpression. **D,** HRAS^G12V^ cotransfection does not alter the relative GDF11 secretion efficiencies of wildtype PCSK5, PCSK5^M452I^, and PCSK5^T288P^. Cells were treated as in (**C**) and measured for GDF11 release by ELISA. For (**A**), (**B**), and (**D**), GDF11 ELISA results are normalized to the GDF11-only condition [gray dashed in (**A**)] and shown as the mean ± SEM of *N* = 4 biological replicates. Differences among +GDF11 groups were analyzed by multiway ANOVA with PCSK5 genotype as a fixed effect. Significant factors were followed up pairwise by Tukey-Kramer post hoc analysis.

We applied the assay to assess enzymatic function of the different PCSK5 alleles. Whereas PCSK5^T288P^ showed no convertase activity, as expected (28), we found that PCSK5^M452I^ activity was consistently detectable but lower than wildtype PCSK5 (**Fig. 2A**, Lanes 5–8). Recognizing that the MCF10DCIS.com genotype is *PCSK5^M452I/+^*, we mixed equal proportions of wildtype and M452I alleles in the cotransfection and repeated the assay. The resulting GDF11 conversion was an average of the two alleles, suggesting additivity in a heterozygous context. We also considered the possibility that a PCSK5^M452I^-specific effect might depend on the inciting HRAS^G12V^ oncogene of MCF10DCIS.com (8). When HRAS^G12V^ was added to the 293T cotransfection and calibrated to the abundance of active prenylated HRAS (54) in MCF10DCIS.com, there was no qualitative change in the measured activity of PCSK5^M452I^ relative to wildtype PCSK5 or PCSK5^T288P^ (**Fig. 2C** and **D**). Contrary to our original hypothesis, the isolated enzymology of PCSK5^M452I^ suggested that it was not hypermorphic but hypomorphic.

### Unmixing PCSK5 alleles in MCF10DCIS.com

The proGDF11 convertase assay estimated enzymatic activity for a transfected PCSK5 allele but did not capture the cellular context of MCF10DCIS.com. We expected that the heterozygous *PCSK5^M452I/+^* genotype of MCF10DCIS.com cells would blur any role for the M452I mutation specifically (**Fig. 1B** and **2B**). A fear of disrupting individual alleles was that heterozygous PCSK5 knockouts would be confounded by clone-to-clone variation given the stem-progenitor characteristics of MCF10DCIS.com (10) and the known impact of GDF11 on stem-cell dynamics (55). We thus adopted a knockout–addback approach that sought to minimize adaptations caused by loss of autocrine PCSK5– GDF11 signaling (**Fig. 3A**). Cells were transduced with lentivirus encoding a destabilization domain-Cas9 (DD-Cas9) fusion (56) and an sgRNA targeting just upstream of the PCSK5 catalytic triad (**Fig. 3B**). MCF10DCIS.com transductants were treated with Shield-1 to stabilize DD-Cas9 (57) plus 2% matrigel to give Cas9 access to the PCSK5 locus (58) for twelve days (**Fig. 3A**). Cells were clonally sorted for Venus co-expression, expanded in low-dose GDF11 to maintain basal signaling during colony recovery, and screened for knockout by sequencing. We moved forward with a *PCSK5^−/−^* clone with morphology and growth characteristics similar to MCF10DCIS.com in 2D culture but with distinct indels yielding premature stop codons for each allele (**Fig. 3B**). This clone was then transduced at limiting multiplicity of infection with a doxycycline-regulated, sgRNA-resistant, and V5-tagged PCSK5 allele (short proteoform; C-terminal tag) and polyclonally selected (**Fig. 3A**). We quantified induction with a V5-containing recombinant standard (59) and detected ∼150,000 copies of each PCSK5 allele after 24 hours of doxycycline (**Fig. 3C**). These stable, inducible, and pure PCSK5 alleles provided a basis for all subsequent experiments involving MCF10DCIS.com.

**Figure 3.**
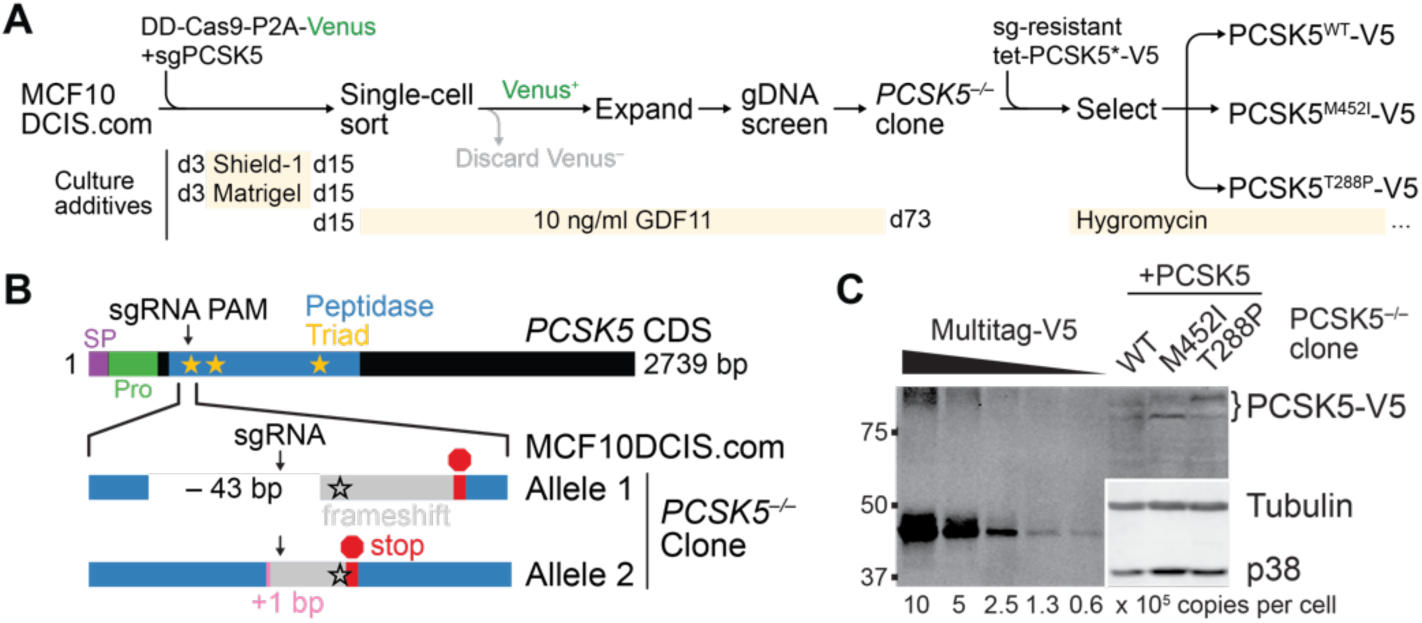
Inducible reconstitution of PCSK5 alleles in PCSK5^−/−^ MCF10DCIS.com cells. **A,** Approach to MCF10DCIS.com engineering. Cells were transduced with a destabilizing domain (DD)-containing Cas9-P2A-Venus (56) and a single-guide RNA targeting Exon 4 of PCSK5 (sgPCSK5). Transduced cells were treated with 200 nM Shield-1 (56) to stabilize Cas9-P2A-Venus and 2% matrigel to promote PCSK5 expression (Fig. 1B) before sorting single Venus-positive cells into 10 ng/ml GDF11 (to aid recovery upon PCSK5 loss) and screening genomic DNA (gDNA) of expanded clones for knockout. One confirmed PCSK5^−/−^ clone was then transduced with sgPCSK5-resistant, tetracycline (tet)-regulated, V5-tagged alleles of PCSK5 and selected polyclonally for hygromycin resistance. **B,** Sequence-confirmed knockout alleles of MCF10DCIS.com clone 3D8. The PCSK5 coding sequence (CDS) is shown with annotations for the signal peptide (SP, purple), proprotein sequence (Pro, green), and peptidase domain (blue) including its catalytic triad (yellow stars). The protospacer adjacent motif (PAM) of sgPCSK5 is just upstream of the first triad amino acid, and deletions (white, Allele 1) or insertions (pink, Allele 2) induce frameshift mutations (gray) removing the first amino acid in the catalytic triad (black outlined stars) and producing premature stop codons (red). **C,** Quantification of reconstituted PCSK5 alleles by calibrating against recombinant V5-containing Multitag protein at the indicated copies per cell (59,68). Cells were treated with 1 µg/ml doxycycline for 24 hours, and total protein from counted cells was immunoblotted for V5 with tubulin and p38 used as loading controls for cells. Copy number estimates are: PCSK5^WT^, 136,000 ± 11,000 copies per cell; PCSK5^M452I^, 164,000 ± 6,000 copies per cell; PCSK5^T288P^, 176,000 ± 15,000 copies per cell (*N* = 4 independent samples).

### Localization of PCSK5 alleles along the distal secretory pathway coincides with auto-maturation

The short proteoform of PCSK5 traffics through the ER and Golgi to secretory vesicles (24,60), and the M452I mutation could affect steady-state localization to these organelles. We leveraged the higher specificity of the V5 antibody (**Fig. 1C** and **3C**; Supplementary Fig. 1) to co-localize reconstituted PCSK5 alleles with markers of the ER [PDI (61)], cis Golgi [GM130 (62)], and trans Golgi [TGN38 (63)]. Using diffraction-limited and adaptively deconvolved image stacks collected by laser-scanning confocal microscopy, we calculated the extent of pixel overlap relative to the overall V5 immunoreactivity per cell. Co-localization with the ER was indistinguishable among PCSK5 alleles (**Fig. 4A** and **B**), but we noted differences in Golgi localization that coincided with whole-cell convertase activity measured earlier (**Fig. 2A**). Compared to wildtype, the inactive T288P mutation accumulated detectably in the cis Golgi and dramatically in the trans Golgi (**Fig. 4C–F**). The hypomorphic M452I mutation, by contrast, was localized similarly to wildtype in the cis Golgi and at an intermediate level between wildtype in T288P in the trans Golgi. These differences were consistent with the extent of proPCSK5 auto-maturation observed by mobility shift in these cells (Supplementary Fig. S3), which occurs after anterograde traffic from the trans Golgi (64). Some V5 was aberrantly immunolocalized to the nucleus, likely from protein that had escaped co-translational import to the ER lumen and undergone nuclear import based on multiple weak localization sequences in the PCSK5 primary structure (Supplementary Fig. S4A). However, the mean nuclear V5 staining was unchanged among PCSK5 alleles (Supplementary Fig. S4B), arguing that M452I is hypomorphic predominantly by inhibiting autoconversion and thus traffic beyond the trans-Golgi network (65).

**Figure 4.**
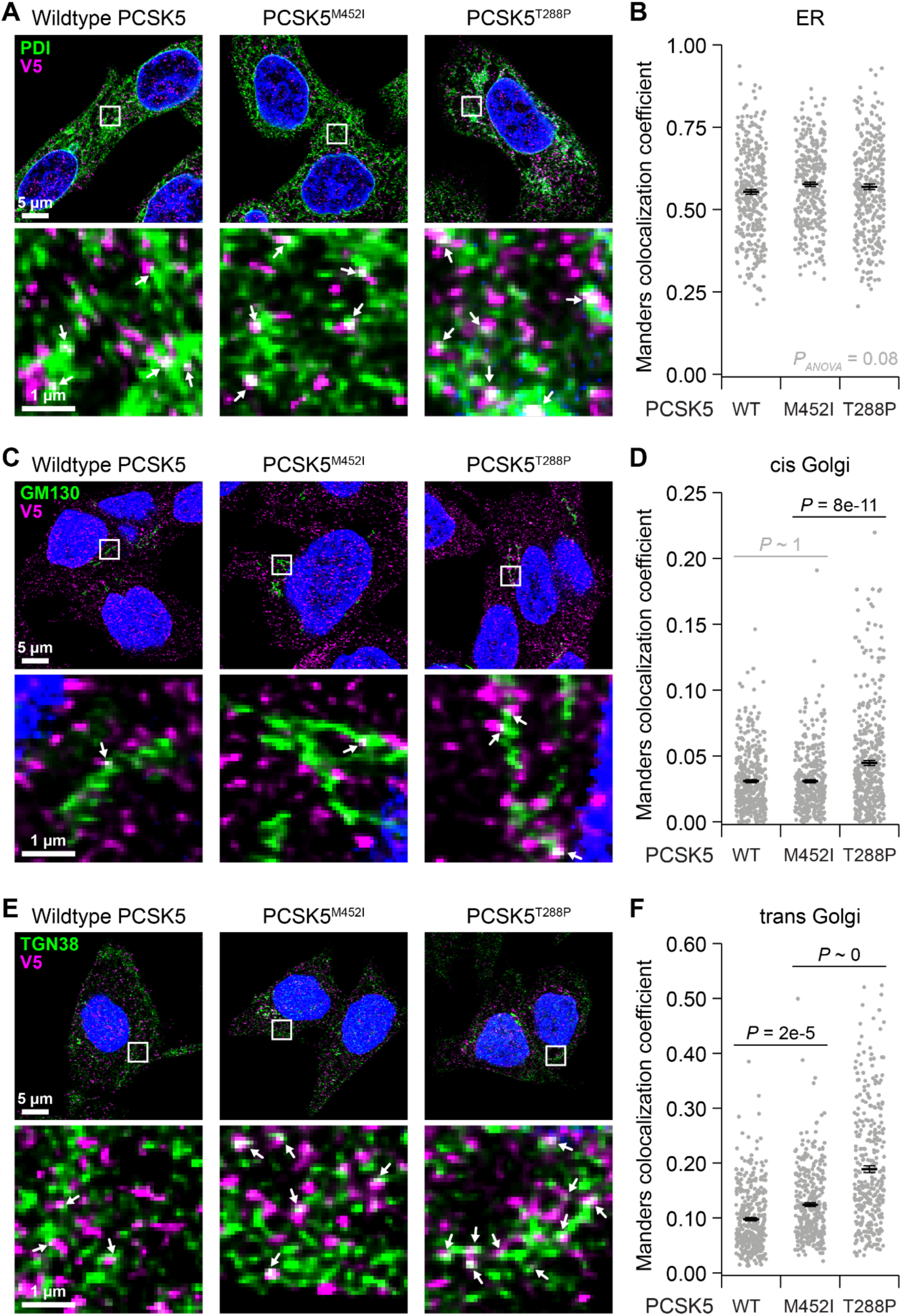
Impaired anterograde transport of catalytically deficient PCSK5 alleles in PCSK5^−/−^ MCF10DCIS.com cells. **A**, Immuno-colocalization of V5 epitope tag (magenta) with the endoplasmic reticulum (ER) marker PDI (green) for the indicated PCSK5 addback allele. **B**, Whole-cell V5–PDI colocalization quantified by Manders colocalization coefficient as a fraction of total per-cell V5 immunoreactivity in *N* = 348 (wildtype PCSK5), 322 (PCSK5^M452I^), and 324 (PCSK5^T288P^) cells. **C**, Immuno-colocalization of V5 epitope tag (magenta) with the cis Golgi marker GM130 (green) for the indicated PCSK5 addback allele. **D**, Whole-cell V5–GM130 colocalization quantified by Manders colocalization coefficient as a fraction of total per-cell V5 immunoreactivity in *N* = 398 (wildtype PCSK5), 373 (PCSK5^M452I^), and 431 (PCSK5^T288P^) cells. **E**, Immuno-colocalization of V5 epitope tag (magenta) with the trans Golgi marker TGN38 (green) for the indicated PCSK5 addback allele. **F,** Whole-cell V5–TGN38 colocalization quantified by Manders colocalization coefficient as a fraction of total per-cell V5 immunoreactivity in *N* = 361 (wildtype PCSK5), 358 (PCSK5^M452I^), and 324 (PCSK5^T288P^) cells. For (**A**), (**C**), and (**E**), cells were immunostained for the indicated targets along with tubulin for cell segmentation, counterstained with DAPI (blue), and imaged on a laser-scanning confocal microscope followed by adaptive image deconvolution. Scale bars are 5 µm (upper) and 1 µm (lower). For (**B**), (**D**), and (**F**), arcsine-transformed coefficients were analyzed by multiway ANOVA with PCSK5 genotype as a fixed effect. Significant factors were followed up pairwise by Tukey-Kramer post hoc analysis.

### PCSK5 deficiency perturbs 3D organization of MCF10DCIS.com spheroids

MCF10-series lines exhibit more phenotypes when overlay cultured with matrigel in 3D (66–68). In MCF10DCIS.com cells, for example, acute knockdown of GDF11 has no discernible effect in 2D culture, but the same perturbation causes 3D spheroids to rupture (28). This result together with the induction of PCSK5 in matrigel (**Fig. 1C**) motivated us to characterize the different PCSK5 addback derivatives of MCF10DCIS.com in 3D.

We digitally segmented (48) hundreds of organoids at multiple time points but did not see gross changes in 3D growth (49) among the PCSK5 alleles (**Fig. 5A**). However, when comparing parental MCF10DCIS.com cells to the originating *PCSK5^−/−^* clone (**Fig. 3A**), there was a notable change in 3D morphology that was quantifiable as spheroid circularity (**Fig. 5B** and **C**). We revisited the three PCSK5 addback lines in 3D and added a high-dose recombinant GDF11 condition to the T288P allele as a positive control for the maximum circularization possible through PCSK5–GDF11 signaling (28). PCSK5^T288P^ cultures were the least spherical, consistent with the null activity of this allele (**Fig. 2A**), and the dramatic increase in circularity with high-dose GDF11 confirmed that they remain responsive to mature ligand (**Fig. 5D** and **E**). Wildtype PCSK5 addback cultures were significantly more circular than PCSK5^T288P^ but did not reach the PCSK5^T288P^ + GDF11 condition, reflecting the MCF10DCIS.com response to endogenous, sub-saturating levels of mature GDF11. Despite earlier biochemical and subcellular evidence of intermediate activity (**Fig. 2A** and **4E** and **F**), we found that circularity of the PCSK5^M452I^ line was not distinguishably different from the null PCSK5^T288P^ allele (**Fig. 5D** and **E**; see Discussion). At the 3D endpoint, we assessed CDH1, TP63, and VIM as markers of epithelial, myoepithelial, and mesenchymal differentiation and did not observe bulk differences among lines without high-dose GDF11 (**Fig. 5F** and **G**). We attribute the induction of VIM in GDF11-stimulated PCSK5^T288P^ cultures to the synergy recognized between TGFβ-superfamily ligands and oncogenic HRAS (69,70). Overall, the multicellular phenotype of PCSK5^M452I^-reconstituted MCF10DCIS.com provides further support that PCSK5^M452I^ is a hypomorph.

**Figure 5.**
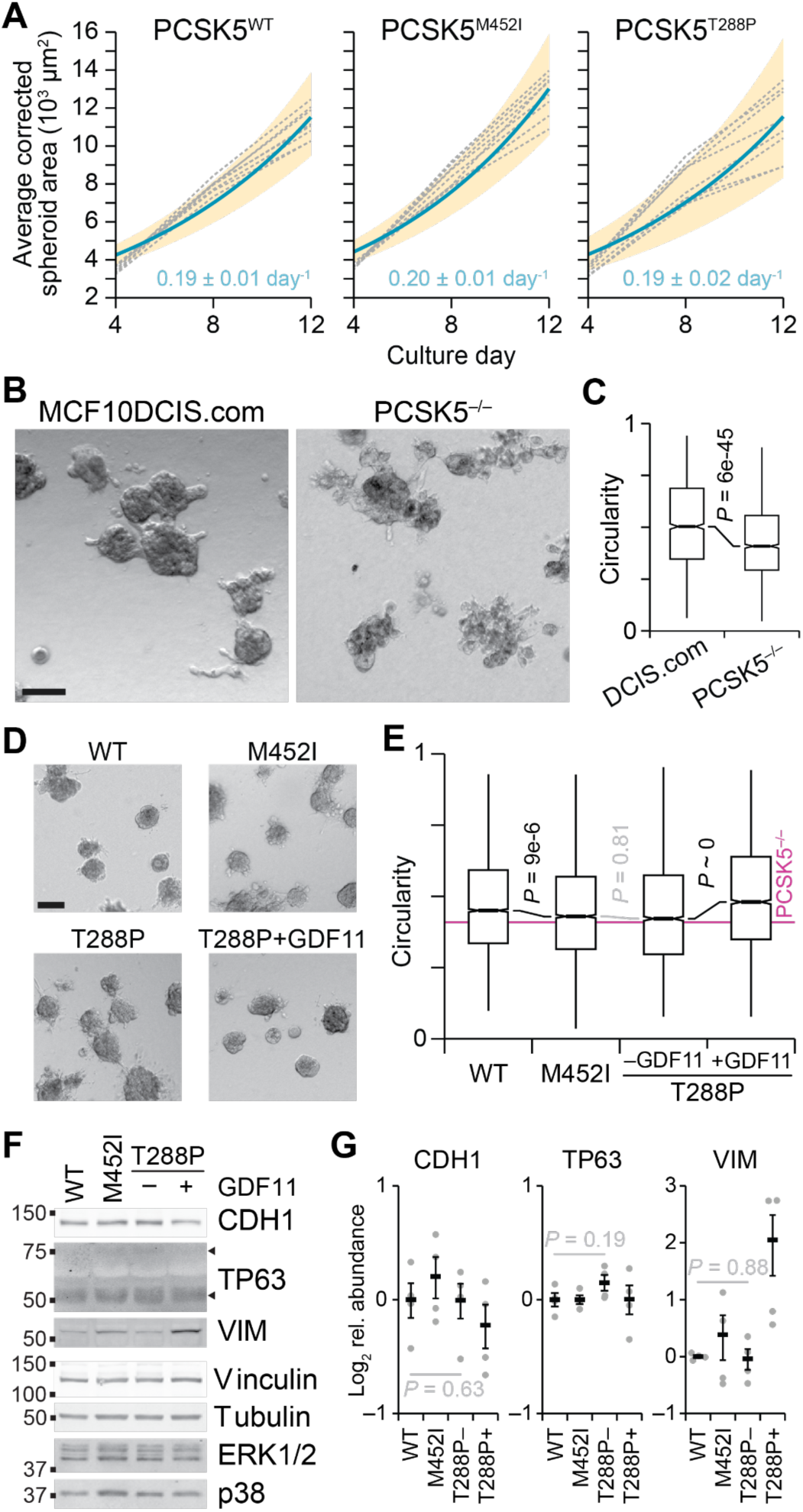
PCSK5 activity promotes rounded multi-cell organization in 3D matrigel cultures of MCF10DCIS.com. **A,** Spheroid growth rates for the indicated PCSK5 addback lines estimated by nonlinear least-squares regression of cross-sectional area (49) at 4, 8, and 12 days from *N* = 8 biological replicates (gray dashed). **B** and **C,** Reduced multi-cell circularity of MCF10DCIS.com upon loss of PCSK5. For (**B**), the scale bar is 100 µm. For (**C**), circularities were calculated from *N* = 1819 (DCIS.com) and 2028 (PCSK5^−/−^) segmented spheroids collected from 4 biological replicates at 16 days. Arcsine-transformed circularities were analyzed by two-sample homoscedastic *t* test. **D** and **E,** Multi-cell circularity of PCSK5^−/−^ cells is restored by wildtype PCSK5 or addition of recombinant GDF11, but not PCSK5^M452I^ or PCSK5^T288P^. For (**D**), the scale bar is 100 µm. For (**E**), circularities were calculated from *N* = 5503 (wildtype PCSK5), 5016 (PCSK5^M452I^), 3495 (PCSK5^T288P^), and 3815 (PCSK5^T288P^+GDF11) segmented spheroids collected from 8 biological replicates at 8 days, and arcsine-transformed circularities were analyzed by multiway ANOVA with PCSK5 genotype as a fixed effect. Significant factors were followed up pairwise by Tukey-Kramer post hoc analysis. **F** and **G,** PCSK5 alleles do not alter the differentiation phenotypes of MCF10DCIS.com cells in 3D matrigel culture. Cultures in (**A**) plus PCSK5^T288P^+GDF11 cultures were lysed and immunoblotted for CDH1, TP63, and VIM with vinculin, tubulin, ERK1/2, and p38 used as loading controls. For (**G**), data from *N* = 4 biological replicates were normalized to the mean of wildtype PCSK5 cultures, and the three unstimulated genotypes were Box-Cox-transformed and compared by multiway ANOVA with PCSK5 genotype as a fixed effect.

### Wildtype PCSK5 promotes the *in vivo* phenotypes of MCF10DCIS.com

MCF10DCIS.com owes its name to the <u>DCIS</u>-like lesions with <u>com</u>edo necrosis formed upon injection in the mammary fat pad (9). To investigate a possible role for PCSK5 in this histopathology, we used intraductal injection (17) to inoculate SCID-beige animals with different PCSK5 addback lines of MCF10DCIS.com. Cells were labeled with luciferase to estimate the surviving inoculum at 2, 7, and 14 days, which together served as an internal reference for outgrowth over the subsequent six weeks after PCSK5 expression was induced with doxycycline at Day 7 (28). We found that wildtype PCSK5 inductions yielded significantly more bioluminescence than PCSK5^M452I^ and PCSK5^T288P^, which were comparable to one another throughout (Supplementary Fig. S5A). Endpoint bioluminescence correlated with estimated tumor volumes upon excision (Supplementary Fig. S5B), verifying that the longitudinal images had captured intraductal growth. Notably, from Day 21 onward, the exponential rate of bioluminescence increase was similar for all genotypes (Supplementary Fig. S5A). Wildtype PCSK5 was distinctive in its growth from Day 14–21, when MCF10DCIS.com xenografts start organizing as DCIS lesions (71) and PCSK5 is presumably increasing from doxycycline induction. The results suggested a transient role for wildtype PCSK5 activity—and, by extension, MCF10DCIS.com-derived GDF11—in intraductal recovery as xenografts, for which PCSK5^M452I^ cannot compensate.

At the endpoint, glands were harvested and carefully assessed by hematoxylin–eosin staining for the per-lesion frequency of various histological phenotypes. We observed no difference in the prevalence of ducts with (micro)papillary, solid, or cribriform growth patterns among PCSK5 alleles, and there was no discernible effect on the frequency of luminal gaps or secretions (Supplementary Fig. S6). Interestingly, we identified two differences that distinguished wildtype PCSK5 or PCSK5^M452I^ from the other alleles. First, comedo necrosis at different length scales was most frequent in glands inoculated with cells harboring wildtype PCSK5 (**Fig. 6A** and **B**). These results are consistent with recent work implicating GDF11 in cell death mediated by hypoxia (72). Second, there were distinctions in the stroma that evolved inside and around MCF10DCIS.com lesions (71). Instances of stromal proliferation in and around the lesions were most frequent with wildtype PCSK5 and least frequent with PCSK5^M452I^ (**Fig. 6C** and **D**). Breast premalignancies with desmoplastic stroma have better clinical outcomes (2), suggesting a role for the stromal response in enforcing the DCIS state. Taken together, these *in vivo* data indicate that PCSK5 activity drives both the *in situ* and comedo phenotypes of MCF10DCIS.com; PCSK5^M452I^ is phenotypically at least as weak as the PCSK5^T288P^ null allele and thus should be impenetrant as a *PCSK5^M452I/+^* heterozygote.

**Figure 6.**
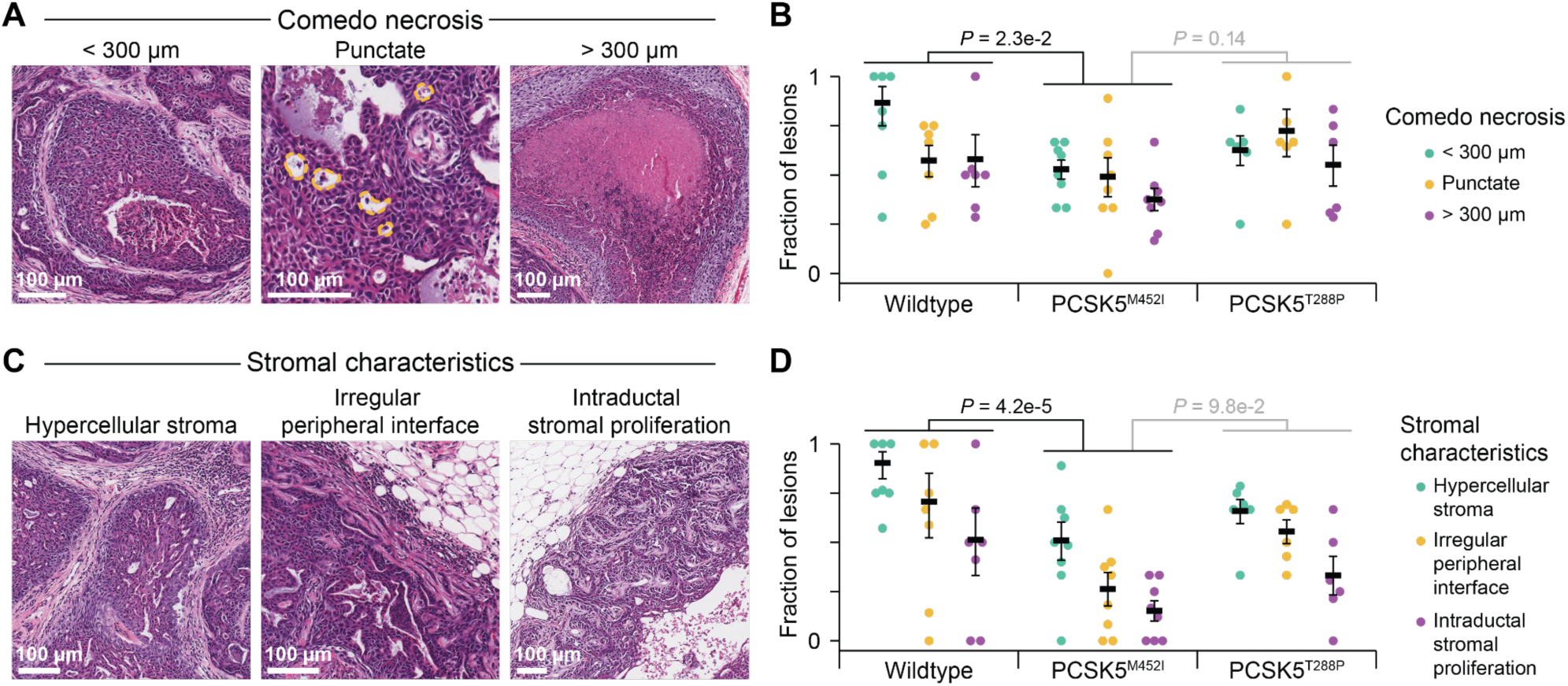
PCSK5 activity promotes comedo and stromal phenotypes in MCF10DCIS.com intraductal xenografts. **A,** Representative hematoxylin–eosin images of wildtype PCSK5 lesions exhibiting small (less than 300 µm; left), punctate (yellow; middle), or large (greater than 300 µm; right) comedo necrosis at 54 days post-injection. **B,** Prevalence of comedo necrosis phenotypes among lesions from *N* = 6–8 animals per PCSK5 genotype. **C,** Representative hematoxylin–eosin images of wildtype PCSK5 lesions exhibiting hypercellular stroma (left), irregularity at the peripheral DCIS-stromal interface (middle), or intraductal stromal proliferation (right) at 54 days post-injection. **D,** Prevalence of stromal phenotypes among lesions from *N* = 6–8 animals per PCSK5 genotype. For (**A**) and (**C**), the scale bar is 100 µm. For (**B**) and (**D**), arcsine-transformed fractions were analyzed by multiway ANOVA with PCSK5 genotype and sub-phenotype as fixed effects. Significant differences by genotype were followed up by Tukey-Kramer post hoc analysis.

## DISCUSSION

This study carefully examines an exclusive PCSK5^M452I^ somatic mutation found in the widely used cell line, MCF10DCIS.com (9,11,23). By comparing to wildtype PCSK5 and catalytically null PCSK5^T288P^ throughout, we exclude PCSK5^M452I^ as a hypermorph and provide strong evidence that it is enzymatically deficient. This deficiency is first reflected as incomplete processing of the inactive zymogen, which retains proPCSK5^M452I^ in the trans Golgi and impedes its anterograde transport to secretory vesicles and the cell surface. Consequently, proGDF11 is unable to mature and accumulates intracellularly (28). Although PCSK5^M452I^ will process GDF11 when both are overexpressed, our results with endogenous GDF11 and minimally reconstituted PCSK5 indicate that PCSK5 activity is rate limiting in MCF10DCIS.com. Observing such PCSK5-dependent phenotypes may require 3D and *in vivo* microenvironments, which trap autocrine factors like GDF11 locally and evolve with time.

Using a new high-sensitivity PCSK5 antibody, we found that endogenous PCSK5 was induced in MCF10-series cells by long-term exposure to matrigel, a form of reconstituted basement membrane. Although the mechanism of induction is unclear, *PCSK5* transcripts are known to increase in breast epithelia upon overexpression of TP53 (73) or knockdown of the BRCA1-interacting protein, BRIP1 (74). In 3D matrigel cultures, TP53 is stabilized sporadically by environmental stresses (19), suggesting a similar route may induce *PCSK5* here. Without matrigel, we were repeatedly unable to knock out PCSK5 in MCF10DCIS.com cells, illustrating the importance of basement membrane for opening the genomic locus. Similar pathways may be active during the early intraductal growth of MCF10DCIS.com *in vivo*, and random monoallelic expression of *PCSK5* (75) might generate mixtures of cells very similar to the addback lines developed here.

The PCSK5-dependent stromal response in xenografts suggests that mature GDF11 from MCF10DCIS.com cells reaches murine fibroblasts. GDF11 induces fibrosis in multiple tissue contexts [reviewed in (76)], and SMAD2 phosphorylation is observed at the tumor-stroma interface in MCF10DCIS.com xenografts (71). Although this signaling was originally attributed to TGFβ, GDF11 also triggers phosphorylation of SMAD2 (28), and their combined action may be important for addressing the stromal compartment in a way that TGFβ alone cannot (77,78).

Our results give reassurance to hundreds of MCF10DCIS.com-themed studies over the past two-and-a-half decades (11). Although PCSK5^M452I^ is an outlier mutation, it appears to do little more than dilute somewhat the cellular convertase available for proGDF11. PCSK5 cleaves additional substrates shared by other convertases but is unique in its activity toward proGDF11 (24,25). Growth of intraductal MCF10DCIS.com xenografts is not affected when GDF11 itself is inducibly knocked down after 14 days (28), implying that a mature GDF11 niche is established very early and dispensable thereafter. Complete loss-of-function mutations documented in breast cancer, such as PCSK5^T288P^, may actually impede breast tumorigenesis if they were to occur during premalignancy.

## Authors’ Contributions

**T. Marohl:** Conceptualization, data curation, formal analysis, funding acquisition, investigation, methodology, validation, visualization, writing – original draft, writing – review & editing. **K.A. Atkins:** Formal analysis, methodology, supervision. **L. Wang:** Data curation, funding acquisition, investigation, project administration, validation, writing – original draft, writing – review & editing. **K.A. Janes:** Conceptualization, formal analysis, funding acquisition, investigation, methodology, project administration, supervision, visualization, writing – original draft, writing – review & editing.

## Acknowledgments

We thank Piotr Przanowski for technical help with the mammary gland harvests, Cameron Wells for providing advance access to OrganoSeg2, and Sameer Bajikar and Daniel Gioeli for critically reviewing this manuscript. This work was supported by grants from the NIH (R01-CA214718 to KAJ, KAA and R50-CA265089 to LW), CBIO experimental funding support from the University of Virginia Comprehensive Cancer Center (PJ04191 to KAJ), a graduate research fellowship from the NSF (2018256910 to TM), a Dean’s Fellowship from the University of Virginia School of Engineering and Applied Science (to TM), and a Distinguished Fellowship from the Commonwealth of Virginia (to TM). Data were partly generated in the University of Virginia Flow Cytometry Core Facility (RRID: SCR_017829), the Research Histology Core Facility (RRID: SCR_025470), the Biorepository and Tissue Research Facility (RRID: SCR_022971), and the Molecular Imaging Core Facility (RRID: SCR_025472), which are partially supported by the UVA Comprehensive Cancer Center support grant (P30-CA044579).

## References

1. Stuart KE, Houssami N, Taylor R, Hayen A, Boyages J. Long-term outcomes of ductal carcinoma in situ of the breast: a systematic review, meta-analysis and meta-regression analysis. BMC Cancer 2015;15:890 doi 10.1186/s12885-015-1904-7.

2. Strand SH, Rivero-Gutierrez B, Houlahan KE, Seoane JA, King LM, Risom T, et al. Molecular classification and biomarkers of clinical outcome in breast ductal carcinoma in situ: Analysis of TBCRC 038 and RAHBT cohorts. Cancer Cell 2022;40(12):1521–36 e7 doi 10.1016/j.ccell.2022.10.021.

3. Cancer Genome Atlas N. Comprehensive molecular portraits of human breast tumours. Nature 2012;490(7418):61–70 doi 10.1038/nature11412.

4. Lin EY, Jones JG, Li P, Zhu L, Whitney KD, Muller WJ, et al. Progression to malignancy in the polyoma middle T oncoprotein mouse breast cancer model provides a reliable model for human diseases. Am J Pathol 2003;163(5):2113–26 doi 10.1016/S0002-9440(10)63568-7.

5. Green JE, Shibata MA, Yoshidome K, Liu ML, Jorcyk C, Anver MR, et al. The C3(1)/SV40 T-antigen transgenic mouse model of mammary cancer: ductal epithelial cell targeting with multistage progression to carcinoma. Oncogene 2000;19(8):1020–7 doi 10.1038/sj.onc.1203280.

6. Hutten SJ, Jonkers J. MIND the translational gap: Preclinical models of ductal carcinoma in situ. Clin Transl Med 2023;13(8):e1376 doi 10.1002/ctm2.1376.

7. Soule HD, Maloney TM, Wolman SR, Peterson WD, Jr., Brenz R, McGrath CM, et al. Isolation and characterization of a spontaneously immortalized human breast epithelial cell line, MCF-10. Cancer Res 1990;50(18):6075–86.

8. Miller FR, Soule HD, Tait L, Pauley RJ, Wolman SR, Dawson PJ, et al. Xenograft model of progressive human proliferative breast disease. J Natl Cancer Inst 1993;85(21):1725–32 doi 10.1093/jnci/85.21.1725.

9. Miller FR, Santner SJ, Tait L, Dawson PJ. MCF10DCIS.com xenograft model of human comedo ductal carcinoma in situ. J Natl Cancer Inst 2000;92(14):1185–6.

10. Hu M, Yao J, Carroll DK, Weremowicz S, Chen H, Carrasco D, et al. Regulation of in situ to invasive breast carcinoma transition. Cancer Cell 2008;13(5):394–406 doi 10.1016/j.ccr.2008.03.007.

11. Puleo J, Polyak K. The MCF10 Model of Breast Tumor Progression. Cancer Res 2021;81(16):4183–5 doi 10.1158/0008-5472.CAN-21-1939.

12. Hutten SJ, de Bruijn R, Lutz C, Badoux M, Eijkman T, Chao X, et al. A living biobank of patient-derived ductal carcinoma in situ mouse-intraductal xenografts identifies risk factors for invasive progression. Cancer Cell 2023;41(5):986–1002 e9 doi 10.1016/j.ccell.2023.04.002.

13. Zhou W, Fong MY, Min Y, Somlo G, Liu L, Palomares MR, et al. Cancer-secreted miR-105 destroys vascular endothelial barriers to promote metastasis. Cancer Cell 2014;25(4):501–15 doi 10.1016/j.ccr.2014.03.007.

14. Possemato R, Marks KM, Shaul YD, Pacold ME, Kim D, Birsoy K, et al. Functional genomics reveal that the serine synthesis pathway is essential in breast cancer. Nature 2011;476(7360):346–50 doi 10.1038/nature10350.

15. Kalaany NY, Sabatini DM. Tumours with PI3K activation are resistant to dietary restriction. Nature 2009;458(7239):725–31 doi 10.1038/nature07782.

16. Wei SC, Fattet L, Tsai JH, Guo Y, Pai VH, Majeski HE, et al. Matrix stiffness drives epithelial-mesenchymal transition and tumour metastasis through a TWIST1-G3BP2 mechanotransduction pathway. Nat Cell Biol 2015;17(5):678–88 doi 10.1038/ncb3157.

17. Behbod F, Kittrell FS, LaMarca H, Edwards D, Kerbawy S, Heestand JC, et al. An intraductal human-in-mouse transplantation model mimics the subtypes of ductal carcinoma in situ. Breast Cancer Res 2009;11(5):R66 doi 10.1186/bcr2358.

18. Fattet L, Jung HY, Matsumoto MW, Aubol BE, Kumar A, Adams JA, et al. Matrix Rigidity Controls Epithelial-Mesenchymal Plasticity and Tumor Metastasis via a Mechanoresponsive EPHA2/LYN Complex. Dev Cell 2020;54(3):302–16 e7 doi 10.1016/j.devcel.2020.05.031.

19. Pereira EJ, Burns JS, Lee CY, Marohl T, Calderon D, Wang L, et al. Sporadic activation of an oxidative stress-dependent NRF2-p53 signaling network in breast epithelial spheroids and premalignancies. Sci Signal 2020;13(627):eaba4200 doi 10.1126/scisignal.aba4200.

20. Frittoli E, Palamidessi A, Iannelli F, Zanardi F, Villa S, Barzaghi L, et al. Tissue fluidification promotes a cGAS-STING cytosolic DNA response in invasive breast cancer. Nat Mater 2023;22(5):644–55 doi 10.1038/s41563-022-01431-x.

21. Peuhu E, Jacquemet G, Scheele C, Isomursu A, Laisne MC, Koskinen LM, et al. MYO10-filopodia support basement membranes at pre-invasive tumor boundaries. Dev Cell 2022;57(20):2350–64 e7 doi 10.1016/j.devcel.2022.09.016.

22. Wagner KU. Know thy cells: commonly used triple-negative human breast cancer cell lines carry mutations in RAS and effectors. Breast Cancer Res 2022;24(1):44 doi 10.1186/s13058-022-01538-8.

23. Maguire SL, Peck B, Wai PT, Campbell J, Barker H, Gulati A, et al. Three-dimensional modelling identifies novel genetic dependencies associated with breast cancer progression in the isogenic MCF10 model. J Pathol 2016;240(3):315–28 doi 10.1002/path.4778.

24. Seidah NG, Prat A. The biology and therapeutic targeting of the proprotein convertases. Nat Rev Drug Discov 2012;11(5):367–83.

25. Essalmani R, Zaid A, Marcinkiewicz J, Chamberland A, Pasquato A, Seidah NG, et al. In vivo functions of the proprotein convertase PC5/6 during mouse development: Gdf11 is a likely substrate. Proc Natl Acad Sci U S A 2008;105(15):5750–5 doi 10.1073/pnas.0709428105.

26. McPherron AC, Lawler AM, Lee SJ. Regulation of anterior/posterior patterning of the axial skeleton by growth/differentiation factor 11. Nat Genet 1999;22(3):260–4 doi 10.1038/10320.

27. Simoni-Nieves A, Gerardo-Ramirez M, Pedraza-Vazquez G, Chavez-Rodriguez L, Bucio L, Souza V, et al. GDF11 Implications in Cancer Biology and Metabolism. Facts and Controversies. Front Oncol 2019;9:1039 doi 10.3389/fonc.2019.01039.

28. Bajikar SS, Wang CC, Borten MA, Pereira EJ, Atkins KA, Janes KA. Tumor-Suppressor Inactivation of GDF11 Occurs by Precursor Sequestration in Triple-Negative Breast Cancer. Dev Cell 2017;43(4):418–35 e13 doi 10.1016/j.devcel.2017.10.027.

29. Tate JG, Bamford S, Jubb HC, Sondka Z, Beare DM, Bindal N, et al. COSMIC: the Catalogue Of Somatic Mutations In Cancer. Nucleic Acids Res 2019;47(D1):D941–D7 doi 10.1093/nar/gky1015.

30. Mei W, Faraj Tabrizi S, Godina C, Lovisa AF, Isaksson K, Jernstrom H, et al. A commonly inherited human PCSK9 germline variant drives breast cancer metastasis via LRP1 receptor. Cell 2025;188(2):371–89 e28 doi 10.1016/j.cell.2024.11.009.

31. Debnath J, Muthuswamy SK, Brugge JS. Morphogenesis and oncogenesis of MCF-10A mammary epithelial acini grown in three-dimensional basement membrane cultures. Methods 2003;30(3):256–68.

32. Janes KA, Wang CC, Holmberg KJ, Cabral K, Brugge JS. Identifying single-cell molecular programs by stochastic profiling. Nat Methods 2010;7(4):311–7 doi 10.1038/nmeth.1442.

33. Cheng J, Novati G, Pan J, Bycroft C, Zemgulyte A, Applebaum T, et al. Accurate proteome-wide missense variant effect prediction with AlphaMissense. Science 2023;381(6664):eadg7492 doi 10.1126/science.adg7492.

34. Tordai H, Torres O, Csepi M, Padanyi R, Lukacs GL, Hegedus T. Analysis of AlphaMissense data in different protein groups and structural context. Sci Data 2024;11(1):495 doi 10.1038/s41597-024-03327-8.

35. Rogers MF, Shihab HA, Gaunt TR, Campbell C. CScape: a tool for predicting oncogenic single-point mutations in the cancer genome. Sci Rep 2017;7(1):11597 doi 10.1038/s41598-017-11746-4.

36. Steinhaus R, Proft S, Schuelke M, Cooper DN, Schwarz JM, Seelow D. MutationTaster2021. Nucleic Acids Res 2021;49(W1):W446–W51 doi 10.1093/nar/gkab266.

37. Tang H, Thomas PD. PANTHER-PSEP: predicting disease-causing genetic variants using position-specific evolutionary preservation. Bioinformatics 2016;32(14):2230–2 doi 10.1093/bioinformatics/btw222.

38. Shihab HA, Gough J, Cooper DN, Stenson PD, Barker GL, Edwards KJ, et al. Predicting the functional, molecular, and phenotypic consequences of amino acid substitutions using hidden Markov models. Hum Mutat 2013;34(1):57–65 doi 10.1002/humu.22225.

39. Chun S, Fay JC. Identification of deleterious mutations within three human genomes. Genome Res 2009;19(9):1553–61 doi 10.1101/gr.092619.109.

40. Li C, Zhi D, Wang K, Liu X. MetaRNN: differentiating rare pathogenic and rare benign missense SNVs and InDels using deep learning. Genome Med 2022;14(1):115 doi 10.1186/s13073-022-01120-z.

41. Sim NL, Kumar P, Hu J, Henikoff S, Schneider G, Ng PC. SIFT web server: predicting effects of amino acid substitutions on proteins. Nucleic Acids Res 2012;40(Web Server issue):W452–7 doi 10.1093/nar/gks539.

42. Vaser R, Adusumalli S, Leng SN, Sikic M, Ng PC. SIFT missense predictions for genomes. Nat Protoc 2016;11(1):1–9 doi 10.1038/nprot.2015.123.

43. Liu X, Li C, Mou C, Dong Y, Tu Y. dbNSFP v4: a comprehensive database of transcript-specific functional predictions and annotations for human nonsynonymous and splice-site SNVs. Genome Med 2020;12(1):103 doi 10.1186/s13073-020-00803-9.

44. Janes KA. An analysis of critical factors for quantitative immunoblotting. Sci Signal 2015;8(371):rs2 doi 10.1126/scisignal.2005966.

45. Wang L, Brugge JS, Janes KA. Intersection of FOXO- and RUNX1-mediated gene expression programs in single breast epithelial cells during morphogenesis and tumor progression. Proc Natl Acad Sci U S A 2011;108(40):E803–12 doi 10.1073/pnas.1103423108.

46. Schnell SA, Staines WA, Wessendorf MW. Reduction of lipofuscin-like autofluorescence in fluorescently labeled tissue. J Histochem Cytochem 1999;47(6):719–30.

47. Stirling DR, Swain-Bowden MJ, Lucas AM, Carpenter AE, Cimini BA, Goodman A. CellProfiler 4: improvements in speed, utility and usability. BMC Bioinformatics 2021;22(1):433 doi 10.1186/s12859-021-04344-9.

48. Borten MA, Bajikar SS, Sasaki N, Clevers H, Janes KA. Automated brightfield morphometry of 3D organoid populations by OrganoSeg. Sci Rep 2018;8(1):5319 doi 10.1038/s41598-017-18815-8.

49. Przanowska RK, Labban N, Przanowski P, Hawes RB, Atkins KA, Showalter SL, et al. Patient-derived response estimates from zero-passage organoids of luminal breast cancer. Breast Cancer Res 2024;26(1):192 doi 10.1186/s13058-024-01931-5.

50. Kosugi S, Hasebe M, Matsumura N, Takashima H, Miyamoto-Sato E, Tomita M, et al. Six classes of nuclear localization signals specific to different binding grooves of importin alpha. J Biol Chem 2009;284(1):478–85 doi 10.1074/jbc.M807017200.

51. Ip MM, Asch BB. Methods in mammary gland biology and breast cancer research. New York: Kluwer Academic/Plenum Publishers; 2000. xvi, 329 p., 4 leaves of col. plates p.

52. Rogers MF, Gaunt TR, Campbell C. CScape-somatic: distinguishing driver and passenger point mutations in the cancer genome. Bioinformatics 2021;37(22):4298 doi 10.1093/bioinformatics/btab654.

53. Ge G, Hopkins DR, Ho WB, Greenspan DS. GDF11 forms a bone morphogenetic protein 1-activated latent complex that can modulate nerve growth factor-induced differentiation of PC12 cells. Mol Cell Biol 2005;25(14):5846–58 doi 10.1128/MCB.25.14.5846-5858.2005.

54. Odeniyide P, Yohe ME, Pollard K, Vaseva AV, Calizo A, Zhang L, et al. Targeting farnesylation as a novel therapeutic approach in HRAS-mutant rhabdomyosarcoma. Oncogene 2022;41(21):2973–83 doi 10.1038/s41388-022-02305-x.

55. Lander AD, Gokoffski KK, Wan FY, Nie Q, Calof AL. Cell lineages and the logic of proliferative control. PLoS Biol 2009;7(1):e15 doi 10.1371/journal.pbio.1000015.

56. Senturk S, Shirole NH, Nowak DG, Corbo V, Pal D, Vaughan A, et al. Rapid and tunable method to temporally control gene editing based on conditional Cas9 stabilization. Nat Commun 2017;8:14370 doi 10.1038/ncomms14370.

57. Banaszynski LA, Chen LC, Maynard-Smith LA, Ooi AG, Wandless TJ. A rapid, reversible, and tunable method to regulate protein function in living cells using synthetic small molecules. Cell 2006;126(5):995–1004 doi 10.1016/j.cell.2006.07.025.

58. Horlbeck MA, Witkowsky LB, Guglielmi B, Replogle JM, Gilbert LA, Villalta JE, et al. Nucleosomes impede Cas9 access to DNA in vivo and in vitro. Elife 2016;5 doi 10.7554/eLife.12677.

59. Lopacinski AB, Sweatt AJ, Smolko CM, Gray-Gaillard E, Borgman CA, Shah M, et al. Modeling the complete kinetics of coxsackievirus B3 reveals human determinants of host-cell feedback. Cell Syst 2021;12(4):304–23 e13 doi 10.1016/j.cels.2021.02.004.

60. De Bie I, Marcinkiewicz M, Malide D, Lazure C, Nakayama K, Bendayan M, et al. The isoforms of proprotein convertase PC5 are sorted to different subcellular compartments. J Cell Biol 1996;135(5):1261–75.

61. Freedman RB. Native disulphide bond formation in protein biosynthesis: evidence for the role of protein disulphide isomerase. Trends Biochem Sci 1984;9(10):438–41 doi 10.1016/0968-0004(84)90152-X.

62. Nakamura N, Rabouille C, Watson R, Nilsson T, Hui N, Slusarewicz P, et al. Characterization of a cis-Golgi matrix protein, GM130. J Cell Biol 1995;131(6 Pt 2):1715–26.

63. Girotti M, Banting G. TGN38-green fluorescent protein hybrid proteins expressed in stably transfected eukaryotic cells provide a tool for the real-time, in vivo study of membrane traffic pathways and suggest a possible role for ratTGN38. J Cell Sci 1996;109 (Pt 12):2915–26 doi 10.1242/jcs.109.12.2915.

64. Mayer G, Hamelin J, Asselin MC, Pasquato A, Marcinkiewicz E, Tang M, et al. The regulated cell surface zymogen activation of the proprotein convertase PC5A directs the processing of its secretory substrates. J Biol Chem 2008;283(4):2373–84 doi 10.1074/jbc.M708763200.

65. Constam DB. Regulation of TGFbeta and related signals by precursor processing. Semin Cell Dev Biol 2014;32:85–97 doi 10.1016/j.semcdb.2014.01.008.

66. Muthuswamy SK, Li D, Lelievre S, Bissell MJ, Brugge JS. ErbB2, but not ErbB1, reinitiates proliferation and induces luminal repopulation in epithelial acini. Nat Cell Biol 2001;3(9):785–92.

67. Debnath J, Mills KR, Collins NL, Reginato MJ, Muthuswamy SK, Brugge JS. The role of apoptosis in creating and maintaining luminal space within normal and oncogene-expressing mammary acini. Cell 2002;111(1):29–40.

68. Wang L, Paudel BB, McKnight RA, Janes KA. Nucleocytoplasmic transport of active HER2 causes fractional escape from the DCIS-like state. Nat Commun 2023;14(1):2110 doi 10.1038/s41467-023-37914-x.

69. Oft M, Peli J, Rudaz C, Schwarz H, Beug H, Reichmann E. TGF-beta1 and Ha-Ras collaborate in modulating the phenotypic plasticity and invasiveness of epithelial tumor cells. Genes Dev 1996;10(19):2462–77 doi 10.1101/gad.10.19.2462.

70. Janda E, Lehmann K, Killisch I, Jechlinger M, Herzig M, Downward J, et al. Ras and TGF[beta] cooperatively regulate epithelial cell plasticity and metastasis: dissection of Ras signaling pathways. J Cell Biol 2002;156(2):299–313 doi 10.1083/jcb.200109037.

71. Tait LR, Pauley RJ, Santner SJ, Heppner GH, Heng HH, Rak JW, et al. Dynamic stromal-epithelial interactions during progression of MCF10DCIS.com xenografts. Int J Cancer 2007;120(10):2127–34 doi 10.1002/ijc.22572.

72. Kraler S, Balbi C, Vdovenko D, Lapikova-Bryhinska T, Camici GG, Liberale L, et al. Circulating GDF11 exacerbates myocardial injury in mice and associates with increased infarct size in humans. Cardiovasc Res 2023;119(17):2729–42 doi 10.1093/cvr/cvad153.

73. Perez CA, Ott J, Mays DJ, Pietenpol JA. p63 consensus DNA-binding site: identification, analysis and application into a p63MH algorithm. Oncogene 2007;26(52):7363–70 doi 10.1038/sj.onc.1210561.

74. Daino K, Imaoka T, Morioka T, Tani S, Iizuka D, Nishimura M, et al. Loss of the BRCA1-interacting helicase BRIP1 results in abnormal mammary acinar morphogenesis. PLoS One 2013;8(9):e74013 doi 10.1371/journal.pone.0074013.

75. Deng Q, Ramskold D, Reinius B, Sandberg R. Single-cell RNA-seq reveals dynamic, random monoallelic gene expression in mammalian cells. Science 2014;343(6167):193–6 doi 10.1126/science.1245316.

76. Frohlich J, Vinciguerra M. Candidate rejuvenating factor GDF11 and tissue fibrosis: friend or foe? Geroscience 2020;42(6):1475–98 doi 10.1007/s11357-020-00279-w.

77. Antebi YE, Linton JM, Klumpe H, Bintu B, Gong M, Su C, et al. Combinatorial Signal Perception in the BMP Pathway. Cell 2017;170(6):1184–96 e24 doi 10.1016/j.cell.2017.08.015.

78. Su CJ, Murugan A, Linton JM, Yeluri A, Bois J, Klumpe H, et al. Ligand-receptor promiscuity enables cellular addressing. Cell Syst 2022;13(5):408–25 e12 doi 10.1016/j.cels.2022.03.001.

