## Supplemental Figures and Tables for "PCSK5^M452I^ is a recessive hypomorph exclusive to MCF10DCIS.com cells"

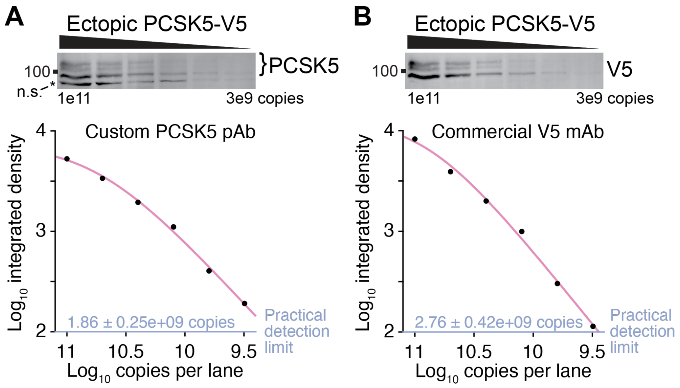


**Supplementary Figure S1.**

Affinity-purified PCSK5 antibody sensitivity and dynamic range.

**A** and **B,** Extract from MDA-MB-231 cells ectopically expressing V5-tagged PCSK5 was calibrated with recombinant V5-containing Multitag protein (1), serially diluted, and immunoblotted for (**A**) PCSK5 and (**B**) V5 as described in the Materials and Methods. Asterisk marks a nonspecific band for the PCSK5 antibody. Bands were quantified by densitometry and fit to a three-parameter logistic equation with regression uncertainties estimated by asymptotic error analysis. Absolute copy-number sensitivities were set to a nominal integrated band intensity of 100 (blue).

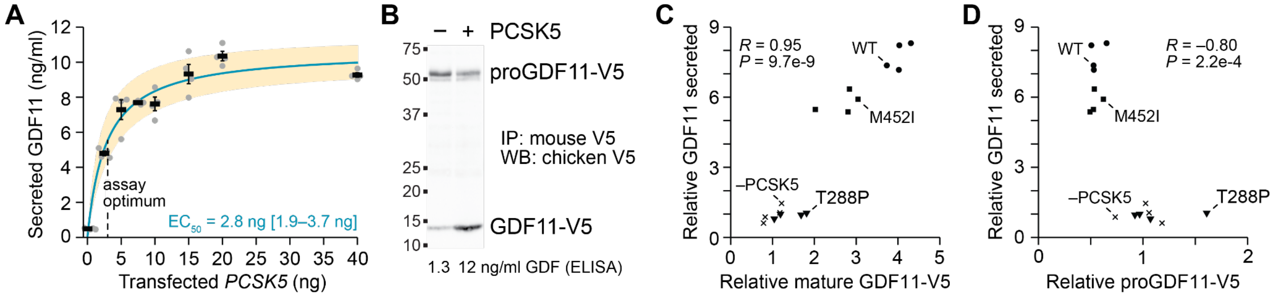


**Supplementary Figure S2.**

Optimization and validation of a proGDF11 convertase assay.

**A,** 293T cells were lipofected with 100 ng of pLX302 GDF11-V5 plasmid plus the indicated mass of pLX304 PCSK5-V5 plasmid in a 24-well plate, and conditioned medium was analyzed for GDF11 release by ELISA. Data are shown as the mean ± SEM of $N = 4$ biological replicates that were fit to a hyperbolic relationship by nonlinear least squares. The half-maximal PCSK5 mass ($EC_{50}$) is shown with 95% confidence interval in brackets, and the dashed line indicates the amount used in the optimized assay. **B,** Conditioned medium from optimized lipofections was collected, immunoprecipitated (IP) with mouse anti-V5, and immunoblotted (WB) with chicken anti-V5 to resolve pro and mature forms of GDF11. ELISA results for these samples are reported underneath the immunoblot. **C** and **D,** Correlation between secreted GDF11 measured by ELISA and (**C**) mature GDF11-V5 or (**D**) proGDF11-V5 measured by immunoblotting. ELISA and immunoblots were normalized to the control lipofections lacking PCSK5 (–PCSK5). Data are from $N = 4$ independent experiments for each PCSK5 genotype and the –PCSK5 control. The Pearson correlation ($R$) was tested for nonzero significance after the Fisher $Z$ transformation.

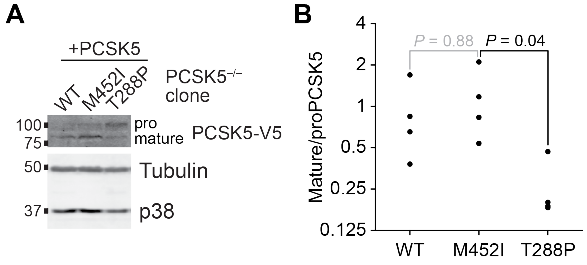


**Supplementary Figure S3.**

Defective maturation of $PCSK5^{T288P}$, but not $PCSK5^{M452I}$, in reconstituted $PCSK5^{-/-}$ MCF10DCIS.com cells.

**A,** Altered relative abundance of pro and mature forms of different PCSK5 alleles. Cells were treated with 1 μg/ml doxycycline for 24 hours, and total protein from counted cells was immunoblotted for V5 with tubulin and p38 used as loading controls. **B,** Densitometry of mature/proPCSK5 for *N* = 4 biological replicates from each genotype. Ratios were analyzed by Kruskal-Wallis test, and significant differences by genotype were followed up by Tukey-Kramer post hoc analysis.

**A**

| Start | PCSK5 sequence | End | cNLS Mapper Mono | cNLS Mapper Bi |
|---|---|---|---|---|
| 52 | RIASKYGFINIGQIGALKDYYHFYHSRTIKRS | 83 | – | 3 |
| 82 | RSVISSRGTHSFISMEPKVEWIQQQVVKKRT | 112 | – | 3.1 |
| 108 | VKKRTKRDYDFSRAQSTYFNDPKWPSMW | 135 | 3 | 3.9 |
| 586 | GKLKEWSLVLYGTSVQPYSPTNEFPKVERF | 615 | – | 3.5 |
| 663 | LKNNTRICVSSCPPGHYHADKKRCRKCAP | 691 | 3.5 | 5.2 |

**B**

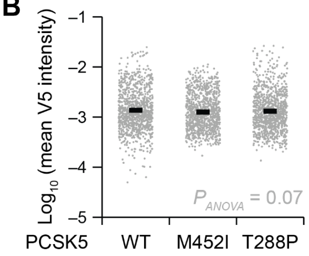


**Supplementary Figure S4.**

Weak nuclear localization signals and observed nuclear localization of PCSK5.

**A,** PCSK5 sequences predicted by cNLS Mapper (2) to be mono- or bi-partite nuclear localization sequences (NLS). Scores of 3–5 predict localization to both the nucleus and the cytoplasm. **B,** Mean nuclear V5 signal intensity in *N* = 1107 (wildtype PCSK5), 1053 ($PCSK5^{M452I}$), and 1079 ($PCSK5^{T288P}$) cells. Log-transformed mean intensities were analyzed by multiway ANOVA with PCSK5 genotype as a fixed effect.

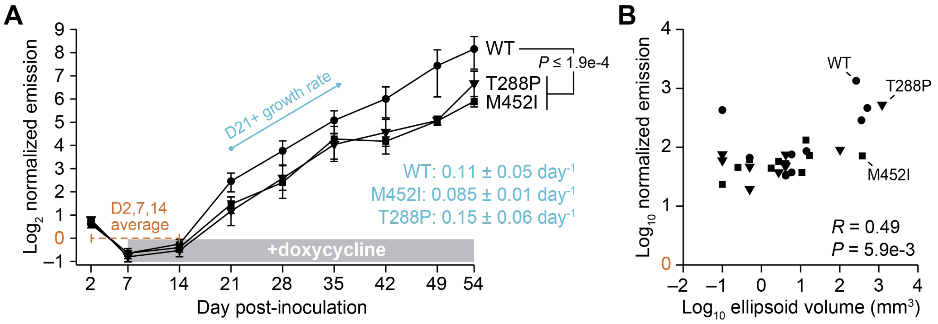


**Supplementary Figure S5.**

Longitudinal tumor bioluminescence and correlation with estimated tumor volume.

**A,** Peak emission was normalized to the Day 2–14 (D2,7,14) average per gland (brown), and data are reported as the mean ± SEM from $N$ = 10 glands per genotype. Doxycycline was added on Day 7. Bioluminescence was compared from Day 21 onward (D21+) by multiway ANOVA with PCSK5 genotype and time point as fixed effects. Significant differences by genotype were followed up by Tukey-Kramer post hoc analysis. Additionally, the D21+ growth rate (blue) was estimated by nonlinear least squares and is shown as the mean ± SEM of the per-day growth rate estimate. **B,** Correlation between peak emission [normalized as in (**A**)] and estimated tumor volume at the study endpoint (Day 54). The Pearson correlation ($R$) was tested for nonzero significance after the Fisher $Z$ transformation.

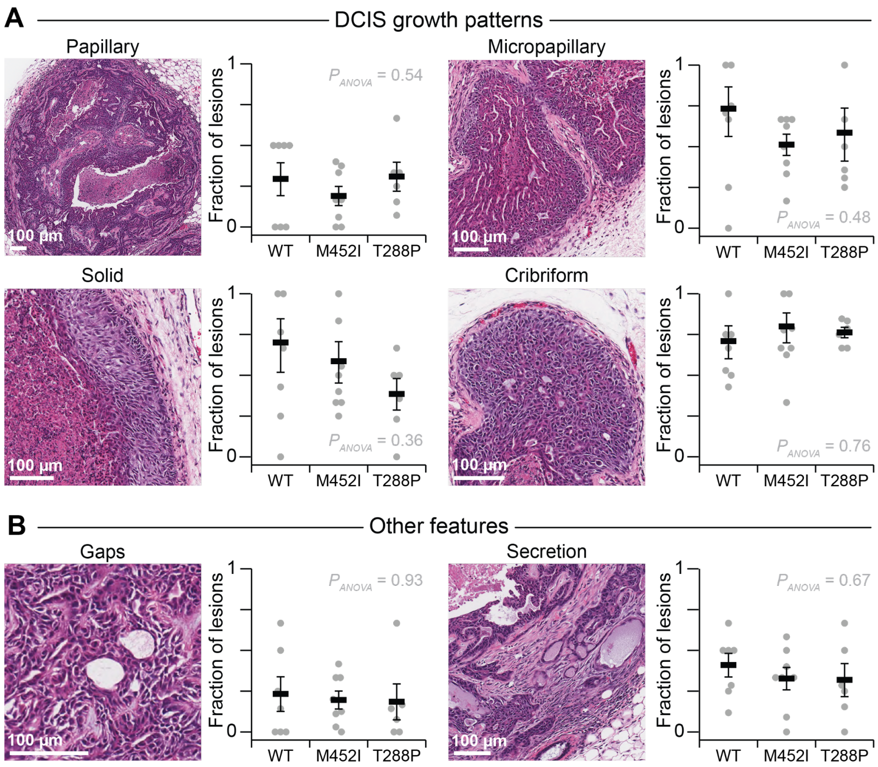


**Supplementary Figure S6.**

Other MCF10DCIS.com histologic phenotypes not detectably altered by PCSK5 activity.

**A,** Papillary (upper left), micropapillary (upper right), solid (lower left), and cribriform (lower right) DCIS growth patterns quantified by lesion prevalence for each genotype. **B,** Gaps (left) and secretions (right) quantified by lesion prevalence for each genotype. For (**A**) and (**B**), hematoxylin–eosin images of wildtype PCSK5 lesions are shown in the left subpanels at 54 days post-injection. The scale bar is 100 µm. In the right subpanels, prevalence among lesions from $N$ = 6–8 animals per PCSK5 genotype was analyzed by multiway ANOVA with PCSK5 genotype as a fixed effect after arcsine transformation of percentages.

**Supplementary Table S1.**

Categorical grouping of PCSK5 mutation predictions.

| | Neutral (N) | Likely Neutral (LN) | Unknown (U) | Likely Damaging (LD) | Damaging (D) |
|---|---|---|---|---|---|
| AlphaMissense | Likely benign | | | | Likely pathogenic |
| CScape | | | | Low-confidence oncogenic | High-confidence oncogenic |
| FATHMM | Tolerated | | | | Damaging |
| LRT | Neutral | | Unreported | | Deleterious |
| Meta-RNN | Tolerated | | | | Damaging |
| MutationAssessor | | Functional—medium | | Non-functional—low | |
| MutationTaster2021 | Benign | | | | Deleterious |
| PANTHER-PSEP | | | | Possibly damaging | Probably damaging |
| SIFT | Tolerated | | | | Damaging |
| SIFT 4G | Tolerated | | | | Damaging |

**Supplementary Table S2.**

Criteria for pathological features scored in MCF10DCIS.com intraductal xenografts.

| Category | Pathologic feature | Description |
|---|---|---|
| DCIS growth patterns | Papillary | MCF10DCIS.com cells arrange around a fibrovascular core |
| | Micropapillary | MCF10DCIS.com cells arrange around a capillary vascular core without a fibrous component to the core |
| | Solid | MCF10DCIS.com cells fill the luminal space without any duct formation in the cellular component |
| | Cribriform | MCF10DCIS.com cells arrange as smaller ducts within the main duct, creating circular or irregular slit-like spaces |
| Comedo necrosis | Large comedo necrosis | Necrotic regions with at least one dimension greater than 300 µm |
| | Small comedo necrosis | Necrotic regions consisting of more than a few cells but with no dimension greater than 300 µm |
| | Punctate necrosis | Necrosis of a single or small number of cells |
| Stromal characteristics | Hypercellular stroma | Increased abundance of periductal stromal cells, immune cells, or both |
| | Irregular peripheral interface | Inter-anastomosing small ducts at the inner surface of the main duct, preventing a smooth DCIS-stroma interface, excluding single cell infiltration |
| | Intraductal stromal proliferation | Stromal cell proliferation that meanders through the ductal space |
| Other | Gaps | Open regions within the duct that lack proliferation of DCIS cells |
| | Secretion | Acellular fluid within the lumens, gaps, or regions of comedo necrosis |

**Supplementary References**


1. Lopacinski AB, Sweatt AJ, Smolko CM, Gray-Gaillard E, Borgman CA, Shah M*, et al.* Modeling the complete kinetics of coxsackievirus B3 reveals human determinants of host-cell feedback. Cell Syst **2021**;12(4):304-23 e13 doi 10.1016/j.cels.2021.02.004.
2. Kosugi S, Hasebe M, Matsumura N, Takashima H, Miyamoto-Sato E, Tomita M*, et al.* Six classes of nuclear localization signals specific to different binding grooves of importin alpha. J Biol Chem **2009**;284(1):478-85 doi 10.1074/jbc.M807017200.
